# Silicon Diode based Flexible and Bioresorbable Optoelectronic Interfaces for Selective Neural Excitation and Inhibition

**DOI:** 10.1101/2022.06.10.495723

**Authors:** Yunxiang Huang, Yuting Cui, Hanjie Deng, Jingjing Wang, Rongqi Hong, Shuhan Hu, Hanqing Hou, Yuanrui Dong, Huachun Wang, Junyu Chen, Lizhu Li, Yang Xie, Pengcheng Sun, Xin Fu, Lan Yin, Wei Xiong, Song-Hai Shi, Minmin Luo, Shirong Wang, Xiaojian Li, Xing Sheng

**Affiliations:** School of Materials Science and Engineering, The Key Laboratory of Advanced Materials of Ministry of Education, State Key Laboratory of New Ceramics and Fine Processing, Center for Flexible Electronics Technology, Tsinghua University, Beijing, 100084, China; Chinese Institute for Brain Research, Beijing, 102206, China; National Institute of Biological Sciences, Beijing, 102206, China; CAS Key Laboratory of Brain Connectome and Manipulation, the Brain Cognition and Brain Disease Institute (BCBDI), Shenzhen Institute of Advanced Technology, Shenzhen-Hong Kong Institute of Brain Science-Shenzhen Fundamental Research Institutions, Shenzhen, 518055, China; School of Life Sciences, Tsinghua University, Beijing, 100084, China; Beijing Advanced Innovation Center for Intelligent Robots and Systems, Beijing Institute of Technology, Beijing, 100084, China; Department of Electronic Engineering, Beijing National Research Center for Information Science and Technology, Center for Flexible Electronics Technology, Tsinghua University, Beijing, 100084, China; IDG/McGovern Institute for Brain Research, Tsinghua University, Beijing, 100084, China

## Abstract

The capability to selectively and precisely modulate neural activities represents a powerful tool for neuroscience research and clinical therapeutics. Traditional electrical stimulations associate with bulky and tethered implants, and optogenetic methods rely on genetic modification for cell targeting. Here, we report an optoelectronic, non-genetic strategy for exciting and inhibiting neural activities, accomplished by bioresorbable, thin-film silicon (Si) diodes. Under illumination, these devices establish polarity-dependent, positive or negative voltages at the semiconductor/solution interface. Such photovoltaic signals enable deterministic depolarization and hyperpolarization of cultured neurons, upregulating and downregulating intracellular calcium dynamics in vitro. Furthermore, flexible, thin-film Si based devices mounted on the nerve tissue selectively activate and silence in vivo activities, both in the peripheral nerve and the brain. Finally, these Si membranes naturally dissolve within the animal body. Such a Si-based material and device platform offers broad potential for biomedical applications.

---

Progresses in neuroscience studies and neurological disease treatments have been significantly impacted by technological advances. Developing the ability to precisely excite and/or inhibit neural activities with high spatiotemporal resolutions represents a key goal for the biological research and clinical translation^1,2,3,4,5,6^. Modulation methods based on electrical^7, 8^ and chemical^9, 10^ cues have been extensively implemented for interrogating the nervous system. Traditional electrical implants accompanied by bulky and invasive constructions could lead to unwanted inflammatory responses^11, 12^, which can be partially mitigated by miniaturized, wirelessly powered circuits^13, 14^. Drug-based modulations with temporal efficacies are commonly lack of real-time precise dosage control, and associated with side effects^15, 16^. More recently, non-invasive or minimally invasive approaches based on alternating electrical field^17^, ultrasound^18^ and magnetism^19^ have also been effectively attempted for neural modulations.

Optogenetic methods, which showcase the high spatiotemporal resolution and minimal invasiveness afforded by light, present a powerful toolbox for precise neural modulations with a high cellular specificity. Various light-sensitive ion channels can be expressed in cell membranes and allow optical-based activation and inhibition at the cellular level as well as in the central and peripheral nervous systems^20, 21^. Nonetheless, the imperative genetic modification for cell targeting impedes their immediate clinical translations. Alternatively, nanoparticles and molecular based photoswitches attached to cell membranes present opto-chemical/mechanical responses and cause non-genetic cell depolarization and hyperpolarization under optical stimulations^22, 23^, while these nanoscale aggregates could accompany cytotoxicity and the uncertainties during the material loading, diffusion and trace in vivo^2^. More recently, light-induced physical stimuli such as photothermal, optoelectronic, photoacoustic and photoelectrochemical effects caused by water^24, 25^, metallic^26, 27^, organic^28,29,30,31,32,33,34,35^, inorganic^36,37,38,39,40^ and graphene^41,42,43^ materials have also been introduced for remote and non-genetic neuromodulations. Photothermal based activation^26, 29, 39, 41^ and inhibition^25, 27, 29^ of neural activities are ascribed to possible mechanisms of the transient high temperature increase for activation and relatively slow small temperature increase for inhibition to block heat-related ion channels^44, 45^. These regulations have to resort to carefully controlled heat strengths and durations in the complex biological environments, to prevent cell dysfunctions and even irreversible damage^46, 47^. On the other hand, optoelectronic processes convert photons to electricity that directly communicates with neurons and other cells like glia, based on photoelectrochemical^36, 38^ and/or photocapacitive effects^31, 32, 37^. While these optoelectronic approaches demonstrate their capabilities for neural excitation in vitro and in vivo, non-genetic neural inhibition functions, which are also crucially important^48^, have been less exploited. Therefore, even though electrically induced cell depolarization and hyperpolarization have been well understood^49, 50^, there is still a pressing need for the development of optoelectronic based, remote techniques to achieve precise neural excitation and inhibition, and we envision such capabilities would enable unprecedented utilities.

Here, we present a remote, non-genetic optoelectronic strategy to realize deterministic neural excitation and inhibition, based on flexible and bioresorbable silicon (Si) membranes. Specifically, thin-film Si based pn diodes create photoinduced positive and negative electric fields, which achieve light-evoked cell depolarization and hyperpolarization and regulate the intracellular calcium dynamics when interfacing with cultured neurons. Furthermore, depending on the polarity of Si diodes, these optoelectronic interfaces also enable selective excitation and inhibition of peripheral and central nervous systems in vivo. Finally, these Si membranes naturally dissolve within the animal body, exhibiting desirable biocompatibilities.

## Results

### Adjustable photovoltaic responses in aqueous solutions

We introduce thin-film, monocrystalline Si based pn diodes to selectively produce photogenerated electrical signals for deterministic neural modulations. Different junctions (p^+^n Si and n^+^p Si) are formed by implanting boron and phosphorous ions into n-type and p-type silicon-on-insulator (SOI) wafers (device layer thickness ∼2 µm), respectively (Fig. S1). Lithographic processes define thin film patterns, and selective etching and transfer printing make freestanding Si films that can heterogeneously integrate onto rigid (like glass) or flexible substrates (see Methods and Fig. S2). Figure 1a presents a released flexible Si film, with its steady-state photovoltage measured by a patch pipette positioned onto the Si surface immersed in aqueous solution (phosphate-buffered saline or PBS, pH = 7.4) (Fig. S3). A red laser beam (peak wavelength 635 nm, ∼2 mm spot size, in continuous mode) is incident on the sample surface and induces intensity-dependent photovoltaic responses in different samples (Fig. 1b and 1c). Both n^+^p and p^+^n diodes dynamically respond to the laser irradiation and present opposite photovoltages in the solution, with photoinduced signals much higher than pure p-type and n-type Si films with the same thickness (∼2 µm). The results can be ascribed to the build-in potential within the pn junctions, which effectively separates photogenerated carriers (electrons and holes) and causes cation or anion accumulation at the Si/solution interface (Fig. 1d). When light intensity varies from ∼0.1 W/cm^2^ to ∼1.8 W/cm^2^, photogenerated voltage ranges between +30 mV and +60 mV for n^+^p Si, and between −30 mV and −60 mV for p^+^n Si. These photovoltages are one order of magnitude lower than the typical open-circuit voltage of a conventional Si photodiode (∼500 mV), which can be explained by the additional potential barrier formed at the Si/solution interface. Nevertheless, these adjustable and dynamic signals can presumably up- and down-regulate cell membrane potentials in the neuronal systems. Like Si photovoltaic cells, these thin-film Si pn junction structures can receive light from both sides and exhibit photo responses across the visible and near-infrared spectra (from 400 nm to 1100 nm). Fig. S4 presents photo responses for these Si diode films under illuminations at different wavelengths (473 nm, 635 nm and 808 nm). The devices generate the largest photovoltages under red light (635 nm), which is consistent with spectral responses of conventional thin-film Si photovoltaic cells with similar thicknesses^51^. In addition, it is noted that here we examine the lightly doped surfaces (p-side for n^+^p Si, and n-side for p^+^n Si) contacting the solution in Fig. 1b and 1c, and we find that the highly doped regions (n^+^-side for n^+^p Si, and p^+^-side for p^+^n Si) also exhibit photogenerated voltages at the Si/solution interface but with much weaker signals (Fig. S5). It is probably attributed to contact barriers and band bending on highly doped Si surfaces that impede carrier accumulation. Therefore, the lightly doped surfaces (p-side for n^+^p Si, and n-side for p^+^n Si) are employed to interface with biological cells and tissues throughout this paper.

**Figure 1.**
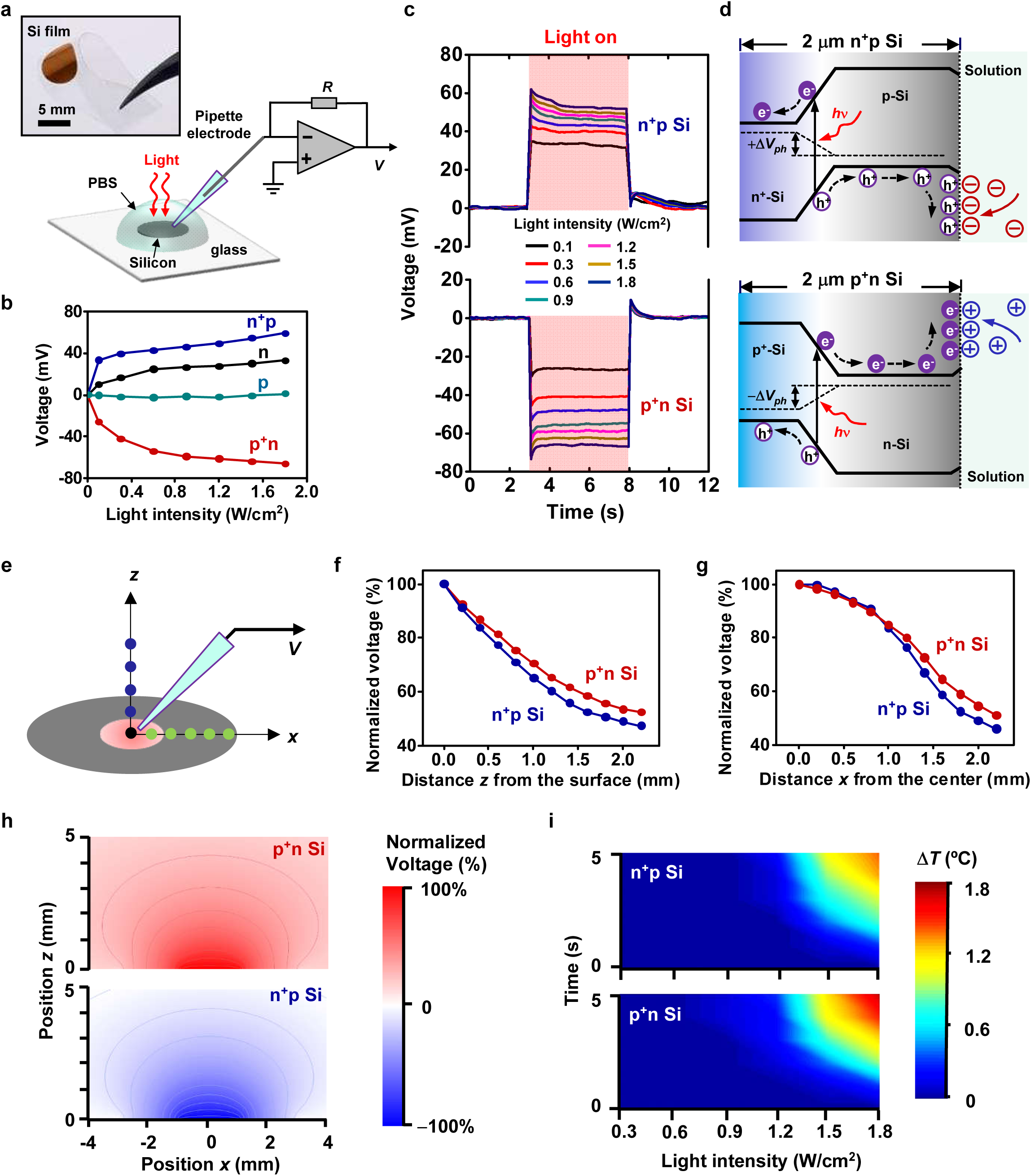
Thin-film Si junctions and their optoelectronic responses in aqueous solutions. **(a)** Image of a freestanding Si film (thickness ∼2 μm, diameter ∼5 mm) transferred onto a flexible polyethylene terephthalate (PET) substrate (up), and scheme of the setup for the photoresponse measurement (down). Light (635 nm laser, ∼2 mm spot size) is incident on the Si film immersed in the PBS solution, with photovoltages recorded with a current-clamp mode. (b) Measured steady-state photovoltages versus light intensity for various types of Si films (p-type, n-type, n^+^p and p^+^n Si junctions) with 5 s continuous illumination. (c) Dynamic photovoltage responses for Si films made of n^+^p and p^+^n junctions, at different light intensities. (d) Operation principles illustrating photogenerated carrier flow (electrons e^−^ and holes h^+^) within the pn junctions as well as charge accumulation (cations and anions) at the Si/solution interface under illumination, for n^+^p and p^+^n Si films, respectively. Dashed lines indicate the quasi-Fermi levels. (e) Scheme for the photoresponse measurement of the 5-mm-diameter Si film at different locations in the solution. (f, g) Measured photovoltage at a function of (f) the vertical distance *z* and (g) the lateral distance *x* from the center of the illuminated spot. (h) Simulated spatial distributions of the electrostatic potential above n^+^p and p^+^n Si films in the solution. The red color presents a positive potential distribution above the p^+^n junction Si, while the blue color presents a negative potential distribution above the n^+^p junction Si. (i) Measured maximum temperature rises on the Si surfaces within the solution, as a function of the illumination duration time and the light intensity.

To evaluate the optoelectronic response of Si films in the aqueous environment, we further quantify the 3D spatial distribution of photovoltage by progressively varying vertical (*z*-axis) and lateral (*x*-axis) distances between the pipette and the illumination center on the Si surface (laser intensity 0.9 W/cm^2^, spot size ∼2 mm) (Fig. 1e). The photovoltaic signals are confined within the illumination spot, and the photovoltage for both n^+^p and p^+^n Si gradually decreases with the increased distances z and x from the center, down to ∼50% of the maximum value at around *z* = 2 mm or *x* = 2 mm (Fig. 1f and 1g). These experimental results are in accordance with calculated photovoltage distributions by finite-element analysis (Fig. 1h). When it comes to resorting photostimulation for neuromodulation, it is also necessary to assess the photothermal effects at the solid/liquid interface. By attaching a thermocouple onto the Si surface, we map the maximum temperature rise occurred on both n^+^p and p^+^n Si, as a function of the illumination duration time and the light intensity (Fig. 1i, Fig. S6). Due to the intrinsic absorption of Si, n^+^p and p^+^n samples present similar photothermal effects. The maximum temperature rise (Δ*T*) can be constrained within ∼0.5 °C when the illumination time is less than 5 s and intensity is lower than 1.2 W/cm^2^, and Δ*T* approaches ∼2 °C when the intensity increases to more than 1.8 W/cm^2^ during 5-s-long illumination. Therefore, carefully controlled illumination conditions can preclude photothermal effects and minimize the heating influence on the neural activities.

### Optoelectronic excitation and inhibition of neural activities in vitro

To investigate the influence of photovoltaic effects on biological systems, we culture rat dorsal root ganglion (DRG) neurons on p^+^n and n^+^p Si films and study photoinduced cell activities using whole-cell patch recordings (Fig. 2a). We first examine the resting membrane potentials (RMPs) of DRGs cultured on p^+^n Si, n^+^p Si and glass (with cell morphologies presented in Fig. S7), and recorded average RMPs for different groups of cells are similar and around −50 mV in dark environment (Fig. 2b). Subsequently, we keep the holding potential at −65 mV and record subthreshold changes of cell membrane potentials induced by illumination with different intensities (Fig. 2c and 2d). Clearly, continuous illumination (duration 5 s) on the p^+^n Si film elevates membrane potentials, resulting in cell depolarization. The shifts of the membrane voltage are consistent with the negative photovoltage generated by the p^+^n Si film, which accumulates cations (Fig. 1b–1d). Conversely, similar illumination conditions on the n^+^p Si film reduce membrane potentials and cause cell hyperpolarization, which can also be explained by the photovoltage induced anion accumulation (Fig. 1b– 1d). Collectively, these remotely controlled, optoelectronic interfaces successfully regulate membrane potentials and cause cell depolarization and hyperpolarization, which previously could only be accomplished by tethered electrodes based on plate capacitors^49, 50^ or pipettes^52^. It is also noted that when the irradiance further increases (> 2 W/cm^2^) in the n^+^p Si film, the membrane potential is inverted from hyperpolarization to depolarization (Fig. 2c, bottom), and eventually leads to the firing of action potentials (APs) (Fig. S8). The reversed intracellular potential and the post-hyperpolarization AP firing after relaxation of the capacitive response are linked with the hyperpolarized potential strength^49^ and the duration of hyperpolarization^54^. This finding prospects possible bidirectional neuromodulation upon the n^+^p Si film through altering light intensity and duration.

**Figure 2.**
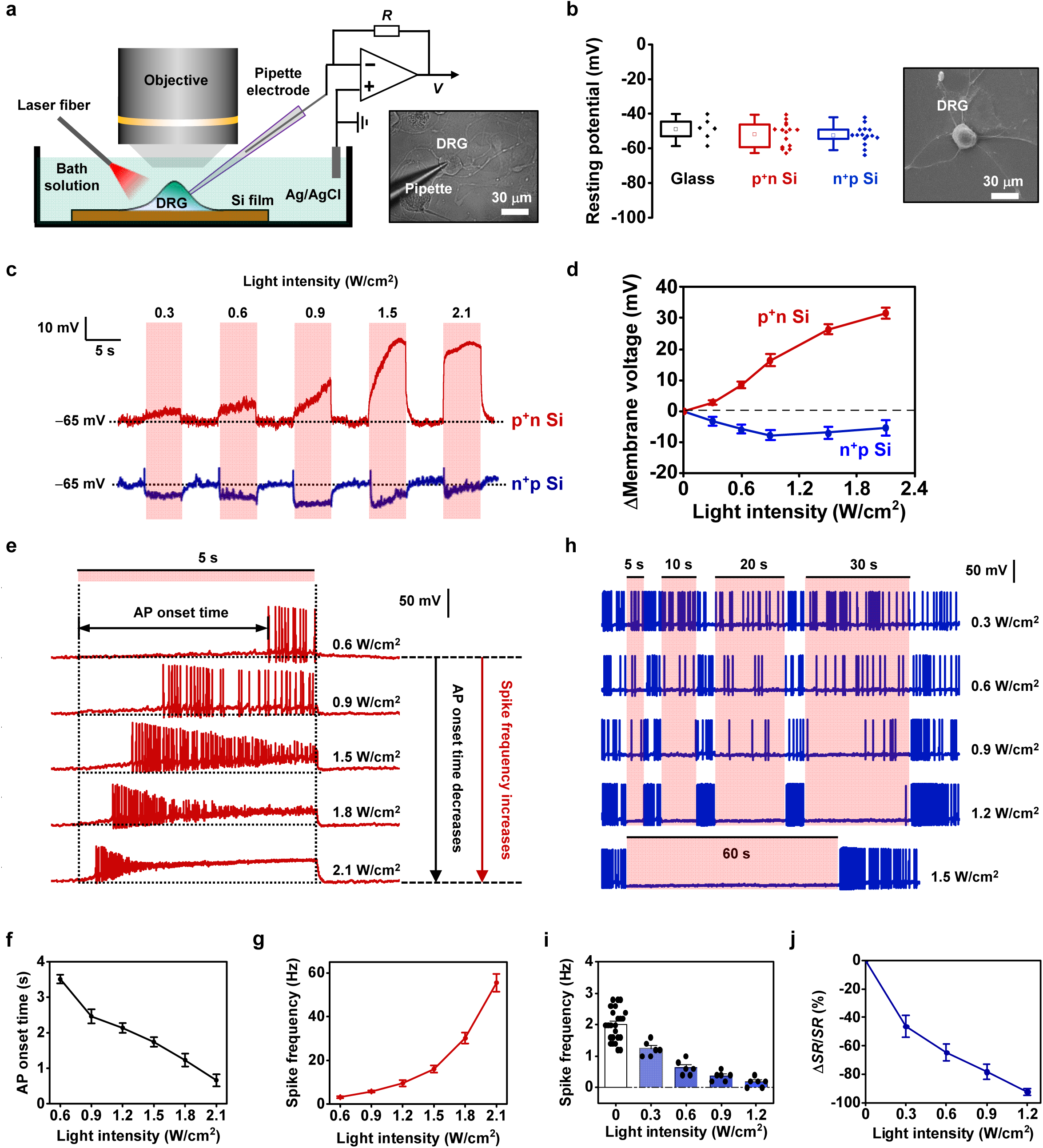
Optoelectronic excitation and inhibition of cell activities *in vitro* with thin-film Si junctions. (a) Left: schematic illustration of the optical neuronal modulation (with a 635 nm laser, ∼1 mm spot size). Right: optical microscopic image of primary rat DRGs cultured on the Si films during patch-clamp recordings. (b) Left: recorded resting membrane potentials for cultured DRGs on glass (*n* = 6 neurons), the p^+^n Si (*n* = 15 neurons) and the n^+^p Si (*n* = 15 neurons). The boxes (25th and 75th percentile), the whiskers (outliers, coefficient 1.5) and the means (square). *p* > 0.2, *t*-test. Right: an SEM image of a DRG neuron on the Si film. (c) Typical traces from current-clamp recordings of the dynamic depolarized (red) and hyperpolarized (blue) membrane voltages generated by differently polarized photoelectric fields during a train of varying light intensities. (d) Statistics of the averaged depolarized and hyperpolarized membrane voltages correlated with the light intensity (*n* = 10 neurons for each). (e) Representative traces of action potentials (APs) elicited by the p^+^n Si in response to different light intensities. (f) Statistics of the onset time for firing the first AP at various light intensities. (g) Statistics of the spike frequency of firing APs from evoking the first one to the last one during the stimulation. Statistics in (f) and (g) are taken from 8 independent neurons with a total of 37 trails. (h) Example traces of inhibiting the excited neurons by the n^+^p Si related to different light intensities at a series of illumination duration time (5 s, 10 s, 20 s and 30 s). Full silence of the excited neuron is maintained for 60 s at an intensity of 1.5 W/cm^2^. The neurons are excited by holding injected currents from pipette electrode. (i) Statistics of spike frequency before and during the stimulation with varying light intensities. Dots present the averaged spike frequency of the 5-s before and duration time of each trail. (j) Statistics of spike rate change (Δ*SR*/*SR*) under different light intensities. Δ*SR*/*SR* is calculated by: [*SF*_(during)_ − *SF*_(before)_]/*SF*_(before)_, where *SF*_(during)_ and *SF*_(before)_ present the averaged spike frequencies (*SF*) during and before the stimulation, respectively. Statistics in (i) and (j) are taken from 6 independent neurons with a total of 24 trails. All data are presented as means ± s.e.m.

We further examine photoinduced cell activities by holding the cell membrane potential at −45 mV. For DRGs on the p^+^n Si film, continuous illumination (5 s) elicits APs (Fig. 2e and Fig. S9). In addition, the AP onset time decreases from 3.5 s to 0.6 s and the spike frequency increases from 3 Hz to 55 Hz when the irradiance increases from 0.6 W/cm^2^ to 2.1 W/cm^2^ (Fig. 2f and 2g). It is noted that the irradiance applied here is three or four magnitudes lower compared to those previously reported Si-based photothermal^39^, photocapacitive^37^ and photoelectrochemical^36^ neuronal excitation of DRGs with pulsed stimulation (pulse duration 0.5–5 ms, intensity 30–60 kW/cm^2^).

On the other hand, Fig. 2c and 2d demonstrate that the illuminated n^+^p Si film can hyperpolarize DRG neurons, which suggests its utility for inhibiting cell activities. In Fig. 2h, DRGs on the n^+^p Si film are activated by continually injecting depolarizing current from pipette, and the cell APs are gradually diminished with increased irradiance, with full suppression obtained at ∼1.5 W/cm^2^ for up to 60 s. The prolonged inhibition is in accordance with the hyperpolarization effect of the illuminated n^+^p Si film (Fig. 2c and Fig. S10). The intensity-dependent inhibition of spike rates is statistically plotted in Fig. 2i and 2j. These Si diode induced photoinhibition effects are distinct from previously reported cases^55, 56^, in terms of their large hyperpolarization potential (∼10 mV), prolonged period (>60 s) and prompt recovery.

These observed cell excitatory and inhibitive responses are primarily induced by photocapacitive effects generated by the Si diode films, since the faradic currents at the the Si/solution interface are negligible (Fig. S11). However, the slowly increased membrane voltage during cell depolarization as well as the existence of the AP onset time indicates that the photoinduced neural response here under continuous illumination has a mechanism distinct from those induced by pulsed light, which usually gives instantaneous AP upon stimulation with delays at millisecond scale^49^. These unusual observations can probably be ascribed to a slowly inactivating potassium current^57, 58^, in accordance with our recorded inward tranmembrane currents (Fig. S12). By contrast, the slowly changing current is not observed when the cell is hyperpolarized (Fig. S13), so the neural inhibition process does not have a delay and occurs almost instantaneously. To further quantitatively understand our experimental results, we adopt a computational model established for extracellular stimulation^49^ and modify it by incorporating the varying transmembrane currents (see Methods and Fig. S14). The simulation results are in a good agreement with our experimental observations (Fig. S15).

As a comparative study, we also investigate DRGs similarly cultured on other substrates including glass, pure p-type Si (p-Si) and n-type Si (n-Si). First, no prominent cell activities are recorded for cells on glass under illumination (light intensity up to 2.1 W/cm^2^, duration up to 5 s) (Fig. S16). Second, similar illumination conditions performed on p-Si and n-Si present negligible influence on DRGs at intensity below 1.0 W/cm^2^ (5 s duration) (Fig. S17 and S18). When the irradiance further increases (1.2−3.2 W/cm^2^), both p-Si and n-Si films produce similar cell depolarization functions, probably due to the enhanced photoactivation in virtue of the semiconductor-solution junction. However, no cell APs are elicited at up to 3.2 W/cm^2^, since the photovoltaic signals from the pure p-Si and n-Si samples are incapable to induce sufficient charges for spike activation. These studies conclude that the tunable, photovoltaic responses of the p^+^n and n^+^p Si diodes represent the key factor that causes neural excitation and inhibition, respectively.

To interrogate neural activities under optoelectronic stimulations at large scale, we perform dynamic calcium (Ca^2+^) imaging of DRGs cultured on Si films (Fig. 3). By loading Fluo-4 AM Ca^2+^ indicator into cells, the recorded intracellular Ca^2+^ fluorescence reveals changes in cell activities that correlates with depolarization and hyperpolarization^37, 41, 50^. For DRGs on the p^+^n Si, fluorescence increases upon illumination (Fig. 3a–3c, Movie S1), and the enhanced irradiance further elevates Ca^2+^ signals (Fig 3d). Apart from the large-sized DRG neurons (∼30 µm), the small-sized glial cells (∼10 µm) neighboring these DRGs also exhibit transient excitation activities responding to illumination (Fig. S19). Conversely, optical stimulations applied for cells cultured on the n^+^p Si (initially activated by α-amino-3-hydroxy-5-methyl-4-isoxazolepropionic acid (AMPA), a specific agonist for the AMPA receptor coupled to ion channels that evoke cell excitation by gating the flow of sodium and calcium ions into the cell) reduce Ca^2+^ fluorescence, indicating the inhibition of cell activities (Fig. 3e–3g, Movie S2). The photo inhibition effect strengthens with increased irradiance (Fig. 3h). Similar to DRGs, glia on the n^+^p Si also exhibit Ca^2+^ signal suppression (Fig. S20). Additionally, these upregulating and downregulating calcium dynamics are clearly observed in cells’ cytosol regions during the photostimulation, which demonstrates intracellular calcium modulation. The excitation and inhibition of Ca^2+^ activities are in good agreement with electrophysiological recordings (Fig. 2), further elucidating the capability of Si diode films in DRG neurons. As a control group, Ca^2+^ loaded DRGs on glass do not experience significant fluorescence changes under illumination (with irradiance up to 3.2 W/cm^2^) (Fig. S21).

**Figure 3.**
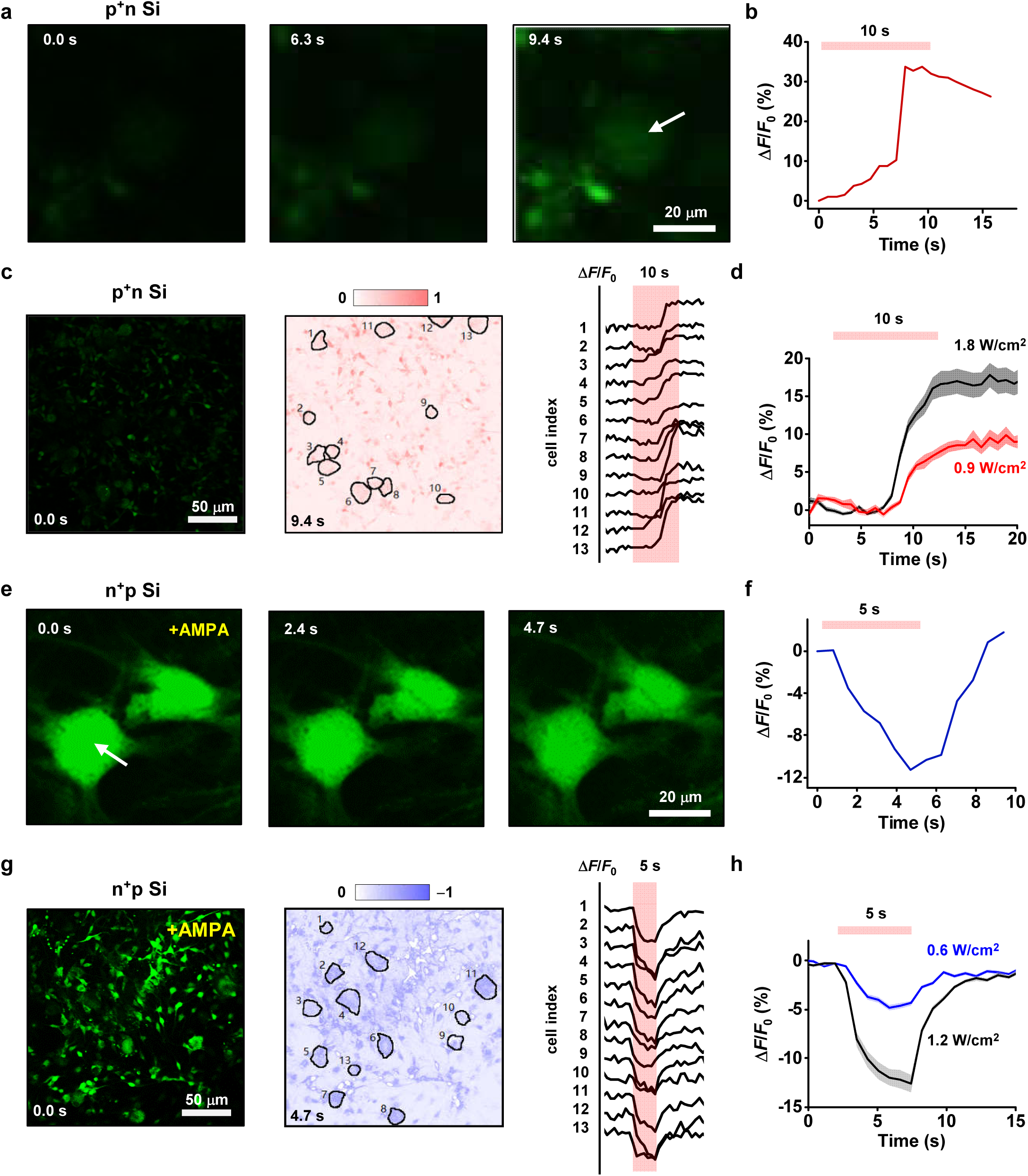
Imaging cellular calcium dynamics optically modulated by thin-film p^+^n and n^+^p Si junctions *in vitro*. (a) Fluorescent images of a typical DRG neuron loaded with Fluo-4 AM cultured on the p^+^n Si, at different time courses under optical stimulation. (b) Calcium signal traces (Δ*F*/*F*_0_) of the arrow-marked DRG neuron (light duration, 10 s; intensity, 2.1 W/cm^2^). (c) Left: Fluorescent image of multiple cells on the p^+^n Si. Middle: Corresponding heat map showing normalized elevated calcium fluorescence before and during the optical activation, with DRG neurons marked by black circles. Red color indicates enhanced fluorescence signal Δ*F*. Right: Calcium signal traces (Δ*F*/*F*_0_) of the marked DRG neurons (light duration, 10 s; intensity, 1.8 W/cm^2^). (d) Statistics of averaged dynamic calcium fluorescence for all marked neurons under different light intensities on the p^+^n Si (duration, 10 s; black curve, 1.8 W/cm^2^; red curve, 0.9 W/cm^2^). (e) Fluorescent images of typical DRG neurons loaded with Fluo-4 AM cultured on the n^+^p Si, at different time courses under optical stimulation. Cells are initially activated by AMPA. (f) Calcium signal traces (Δ*F*/*F*_0_) of the arrow-marked DRG neuron (light duration, 6 s; intensity, 1.5 W/cm^2^). (g) Left: Fluorescent image of multiple cells on the n^+^p Si. Bright calcium fluorescence observed in the region presents activated cellular activities initially evoked by AMPA. Middle: Corresponding heat map showing normalized decreased calcium fluorescence before and during the optical activation, with DRG neurons marked by black circles. Blue color indicates declined fluorescence signal Δ*F*. Right: Calcium signal traces (Δ*F*/*F*_0_) of the marked DRG neurons (light duration, 5 s; intensity, 1.2 W/cm^2^). (h) Statistics of averaged dynamic calcium fluorescence for all marked neurons under different light intensities on the n^+^p Si (duration, 5 s; black curve, 1.2 W/cm^2^; blue curve, 0.6 W/cm^2^). Statistics in (d) and (h) are calculated from calcium images taken from 10 regions of 4 different cultures (*n* > 30 neurons for each condition). All data are presented as means ± s.e.m.

### Optoelectronic excitation and inhibition of peripheral neural activities in vivo

The Si based membrane structures integrated on thin, flexible substrates allow their conformal contact with biological tissues and organs, enabling versatile functions for biological sensing and modulation^59^. Here we apply the thin-film Si diodes (n^+^p and p^+^n) in the peripheral nervous system (PNS), and in particular, evaluate their capabilities in optical excitation and inhibition of the sciatic nerve (Fig. 4). Conventionally, electrical modulations in PNS rely on tethered extraneural electrodes and circuit systems, which resort to low-frequency stimulations for exciting APs and high-frequency pulses for inhibiting AP transmission^8^. Our in vitro results on cultured DRGs (Fig. 2 and 3) implicate that these biocompatible, Si-based diode films could permit wireless, non-genetic optoelectronic modulation in PNS in vivo. We examine the optical modulation effects of Si diode films on the sciatic nerve of wild-type mice without genetic modification (Fig. 4, Fig. S22). Thin-film Si diodes are transferred onto flexible polyethylene terephthalate (PET) substrates and conformally attached on the exposed sciatic nerve. Here the Si surface is decorated by a thin layer of gold nanoparticles, to minimize the silicon-tissue impedance and enhance the efficacy of optoelectronic stimulation in vivo^7, 37, 38^. A laser beam (635 nm) is remotely incident on the Si film. Compound muscle action potentials (CMAPs) and displacements of hindlimb motion track are captured by a recording electrode and a camera, respectively. Pulsed illuminations incident on the p^+^n Si film evoke CMAPs and cause hindlimb lifting (Fig. 4b, Fig. S23, Movie S3). The behavior can be explained by the fact that the optically excited p^+^n Si film induces more positive charges (+Δ*V*) in the nerve fiber, causing cell depolarization and evoking CMAPs (Fig. 4c). By contrast, the n^+^p Si film does not elicit hindlimb activities under similar optical stimulations (Fig. S24), precluding photothermal effects. Additionally, further increased irradiance (> 1.5 W/cm^2^) reduces CMAPs and displacements (Fig. 4d and 4e), possibly due to the refractory period that discourages hindlimb movements under intense stimulation.

**Figure 4.**
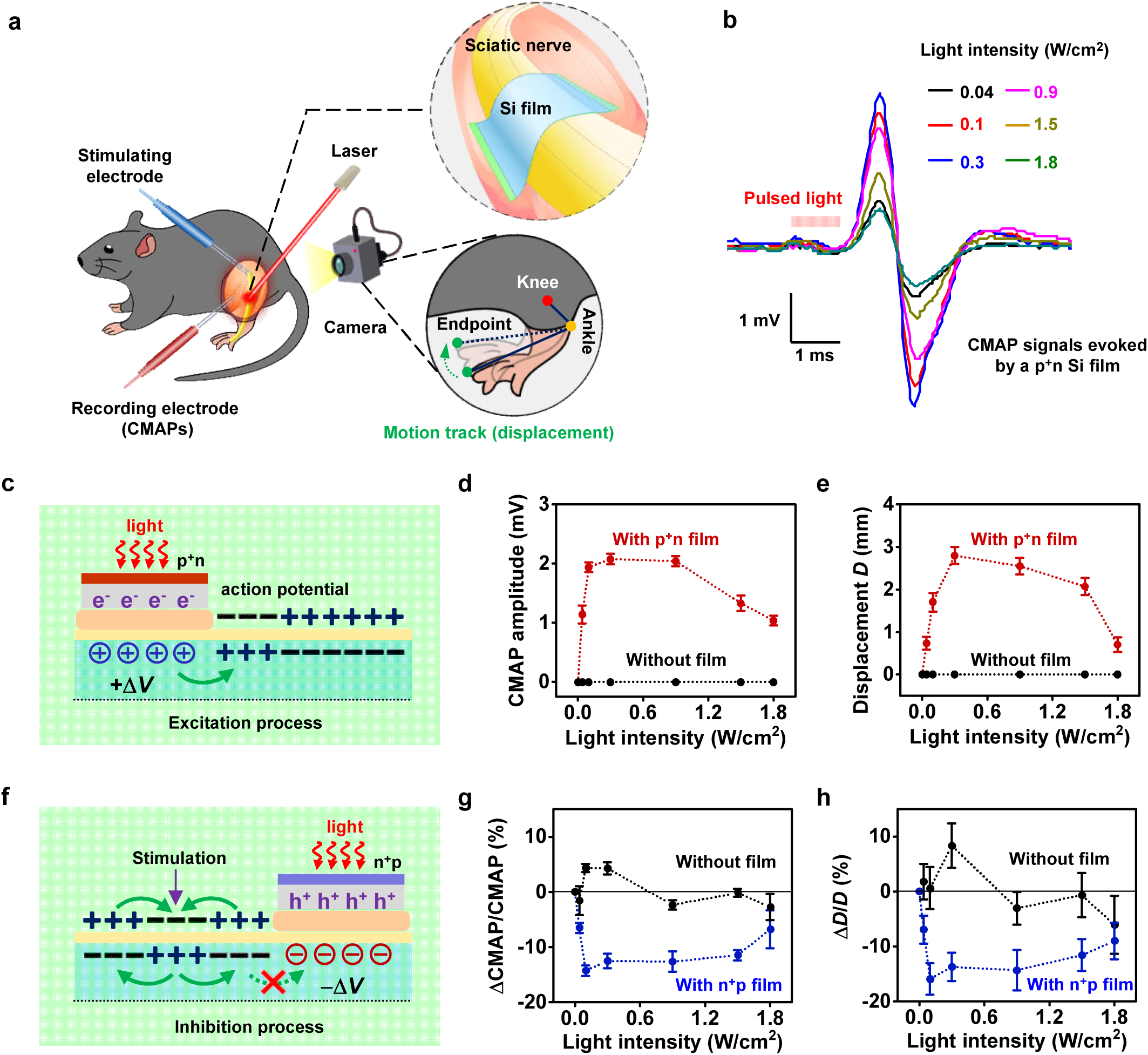
Optoelectronic excitation and inhibition of peripheral neural activities *in vivo* with p^+^n and n^+^p Si films, respectively. (a) Cartoon diagram illustrating the modulation of mice’s hindlimb movement, with light illuminating Si films attached on the sciatic nerve. Compound muscle action potentials (CMAPs) are recorded by EMG recording electrode inserted into the hindlimb-related muscles. Motion tracks are captured by a camera. An external stimulating electrode placed in the proximal position of the nerve evokes hindlimb movements. (b) Typical CMAPs evoked by illuminating the p^+^n Si with varying light intensities (1-ms pulse duration) on one hindlimb. (c) Principle of the excitation process, showing that the electric field generated by the p^+^n Si induces cation accumulation within the neuron and causes cell depolarization. (d, e) Statistics of recorded (d) CMAP amplitudes and (e) limb displacements *D* with and without the p^+^n Si film at varying light intensities (1-ms pulse duration, 0.4 Hz, 9 pulses). Each illumination is repeated more than 2 times on three mice. (f) Principle of the inhibition process, showing that the electric field generated by the n^+^p Si induces anion accumulation within the neuron, causes cell hyperpolarization and block the signal transduction from the proximal position. (g, h) Statistics of normalized inhibition rate (g) ΔCAMP/CAMP and (h) displacements Δ*D*/*D* with and without the n^+^p Si film at varying light intensities (10-s continuous stimulation) under external electrical stimulation. Each illumination is repeated more than 2 times and each time inhibited 40 trails (40-time movements) on three mice. All data are presented as means ± s.e.m.

The above results on the peripheral nerve activation by illuminating the p^+^n Si film have been similarly reported in previous works based on other optoelectronic materials and devices^33^. A more prominent feature of the Si diode based film is that an inverted structure (the n^+^p Si diode) can alternatively achieve photoinduced inhibition. Illustrated in Fig. 4f, the photogenerated negative charges (−Δ*V*) in the nerve can block the positive-charge signal transmission from proximal to distal positions, suppressing CMAPs. In this experiment, an external stimulating electrode regularly evokes hindlimb movements at the proximal nerve position (electrical pulse duration 1 ms, frequency 4 Hz), and continuous illumination (duration 10 s) is incident on the n^+^p Si film attached on the nerve. At irradiance around 0.1–1.5 W/cm^2^, applied optical stimulations reduce electrically evoked CMAPs and displacements, relatively by 10%−15% compared to the averaged data collected before illumination (Fig. 4g and 4h, Fig. S25, Movie S4). By contrast, illumination directly imposed on the nerve (without Si films) does not cause significant inhibition effects. Additionally, a p^+^n Si film applied on the nerve does not exhibit inhibition functions under similar continuous illumination (Fig. S26), further precluding photothermal effects. Unlike results obtained on cultured DRG neurons in vitro (Fig. 2h), here the illuminated n^+^p Si film does not completely suppress in vivo neural activities. These differences follow from multiple reasons: (1) The sciatic nerve comprises bundled fibers naturally packed by multi-layered tissues including endoneurium, perineurium and epineurium^8^, making the Si film not in direct contact with the axons; (2) The Si film mounts on one side of the fiber, only partially inhibiting neural activities; (3) The laser spot only illuminates a small area and does not completely cover the circumference of the nerve fiber; (4) Both excitatory and inhibitory neurons exist within the peripheral nerve system, further complicates in vivo behaviours. All these considerations limit the efficacy of photoinduced inhibition in vivo. In addition to carefully optimized illumination power and time, other strategies to improve the inhibition performance include the implementation of cuff-style electrodes to conformally wrap around the nerve^33^, as well as waveguide structures to distribute uniform illumination around the tissue. Nevertheless, the thin-film Si diodes showcase the capability for selective activation and inhibition of the PNS.

### Optoelectronic excitation and inhibition of cortical neural activities in vivo

Besides the PNS, thin-film Si diodes transferred onto flexible substrates can also mount on the cerebral cortex of animals and modulate in vivo brain activities via their photovoltaic signals (Fig. 5). Schematically illustrated in Fig. 5a and b, Au-coated Si films attach on the somatosensory cortex of a mouse brain. A laser beam (473 nm) is incident on the Si surface, and a linear microelectrode array slantwise inserts into the tissue (with a depth of ∼800 μm from the pia mater) underneath the film to record extracellular activities. The spontaneous and stimulated spike-like events are sorted, showing typical extracellular electrophysiological recording (Fig. 5c). Fig. 5d shows representative electrophysiological signals by illuminating the p^+^n Si film on the cortex (duration 1 s) and corresponding quantified raster plots from the raw data. The heat map from 8 channels depicts spike firing changes inside the layers of the cortex, caused by the photostimulation of the p^+^n Si film (Fig. 5e). These results reveal strong stimulation responses occurred in superficial neurons. These photoactivation results are in accordance with the literature^37^, but a more striking finding is that photoinhibition can be obtained by reversing the diode polarity. Fig. 5f shows typical data recorded for the case with the n^+^p Si film, with the brain activity initially activated by AMPA. The illuminated n^+^p Si film significantly suppress the pre-evoked cortical activities (Fig. 5g). For comparison, results are also recorded for animals with pure p-Si films mounted on the cortex under similar illumination conditions, in order to preclude photothermal effects (Figs. S27 and S28). Statistical data are summarized and presented in Fig. 5h–5m. Clearly, illumination on the p^+^n Si film triggers neuron responses, leading to a significantly enhanced spike frequency compared to the spontaneous state. On the other hand, the pre-evoked neural activities by AMPA can be suppressed by the optically stimulated n^+^p Si film on the cortex. These excitation and inhibition processes can sustain for more than 10 s by introducing 1-s illumination. On the contrary, illuminations imposed on the p-Si film have no significant influences on brain activities. These findings elucidate that the photoinduced, junction dependent electrical fields, rather than the photothermal effect, selectively elicit excited and inhibited neural responses.

**Figure 5.**
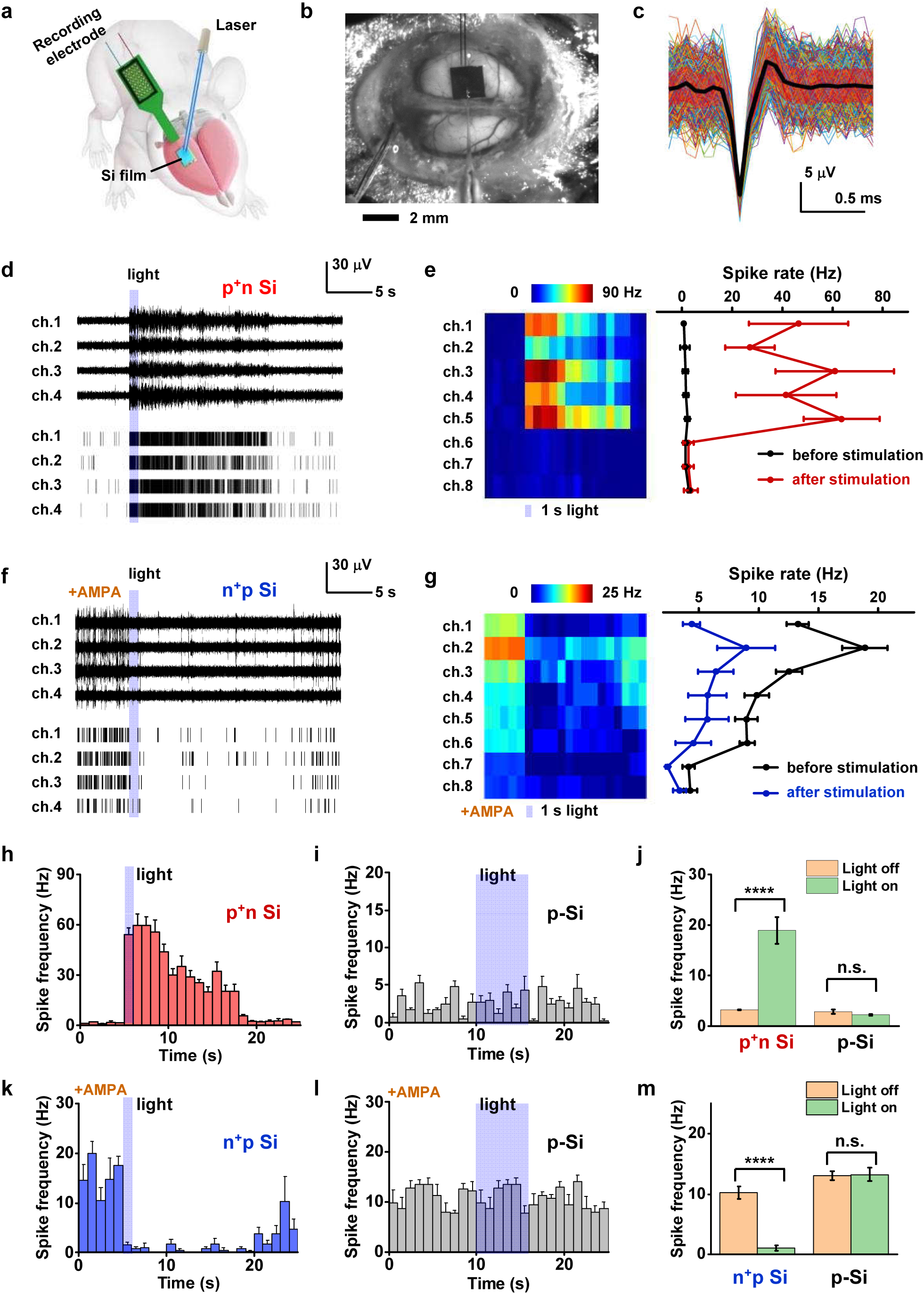
Optoelectronic excitation and inhibition of brain cortex activities *in vivo* with p^+^n and n^+^p Si films, respectively. (a) Illustration and (b) Photograph of the experiment. A multichannel recording probe is guided into mouse brain to sample extracellular activities by illuminating the Si films attached on the cortex with a 473 nm laser. (c) A mean neuron-firing waveform (black) with standard deviations (red shaded area) based on 200 individual waveforms from the spontaneous and stimulation-evoked events in one representative photostimulation experiment on one mouse. (d) and (f) Example traces of recorded raw signals (top) and the corresponding spike raster plots (bottom) from four channels in a single trial, under stimulation by p^+^n and n^+^p Si films, respectively. (e) and (g) Heat maps of channels from 1 to 8 (left) and the mean spontaneous and evoked mean neural response for the same channels in the heat map (right), under 1 s light stimulation (marked below the map) by p^+^n and n^+^p Si films, respectively. In (f) and (g), AMPA is applied to activate neural activities before the illumination. (h, i) Averaged spike frequency from the four channels recorded from animals with (h) the p^+^n Si and (i) the p-Si (intensity 0.06 W/cm^2^, duration 1 s for p^+^n Si and 5 s for p-Si). (j) Statistics of the spike frequency for p^+^n Si and p-Si before and after optical stimulation, at varying light intensity (0–0.32 W/cm^2^) and pulse duration (1–5 s), averaged among *n* = 144 trails (before 10 s and during-after 10 s; **** *p* < 0.0001, *t*-test). (k, l) Averaged spike frequency of the four channels recorded from animals (initially activated by AMPA) with (k) the n^+^p Si and (l) the p-Si (intensity 0.03 W/cm^2^, duration 1 s for n^+^p Si and 5 s for p-Si). (m) Statistics of the spike frequency for n^+^p Si and p-Si before and after optical stimulation, at varying light intensity (0–0.03 W/cm^2^) and pulse duration (1–5 s), averaged among *n* = 192 trails (before 10 s and during-after 10 s; **** *p* < 0.0001, *t*-test). *n* = 3 mice for each experiment. All data are presented as means ± s.e.m.

### Biodegradation of thin-film Si diodes

Compared to other optoelectronic materials based on III–V (GaAs, GaN, etc.), organic semiconductors^28, 31,32,33,34,35^ and inorganic nanocrystals^40^, another unique characteristic for thin-film Si based devices is their compatibility and even degradability within biological systems^60, 61^. Here we study chronic properties and dissolution processes of Si films in vitro and in vivo (Fig. 6). p^+^n and n^+^p Si films are immersed in aqueous solutions (0.1 M PBS, pH = 7.4, 37 °C) for up to 5 months, and evolutions of film thicknesses and photo responses are measured and presented in Fig. 6a and 6b, respectively. Averaged dissolution rates are ∼2 nm/day for p^+^n Si and ∼3 nm/day for n^+^p Si, consistent with previous reports^62^. Along with the film dissolution, their photovoltages drop more dramatically, to ∼40% of the original values after 5 months in PBS. In the meantime, we wrap Si films around mice’s sciatic nerve (Fig. 6c) and attach Si films on mice’s brain cortex (Fig. 6d), evaluating their in vivo degradation. These Si films completely disappear after 5 months, showing a faster degradation compared to in vitro results. Histological images of the sciatic nerve show minimal damage to the tissue and negligible immune response associated to the implantation (Fig. 6e). Elemental analyses are performed for the brain tissue with Si implants, and minimal accumulation of dissolved Si is observed (Fig. 6f). Moreover, biochemical tests of animal blood also indicate no physiological abnormalities after implantation (Fig. S29). The natural dissolution of Si films with desirable biocompatibilities in the animal body eliminates the requirements for secondary surgery, showing promises for certain medical applications.

**Figure 6.**
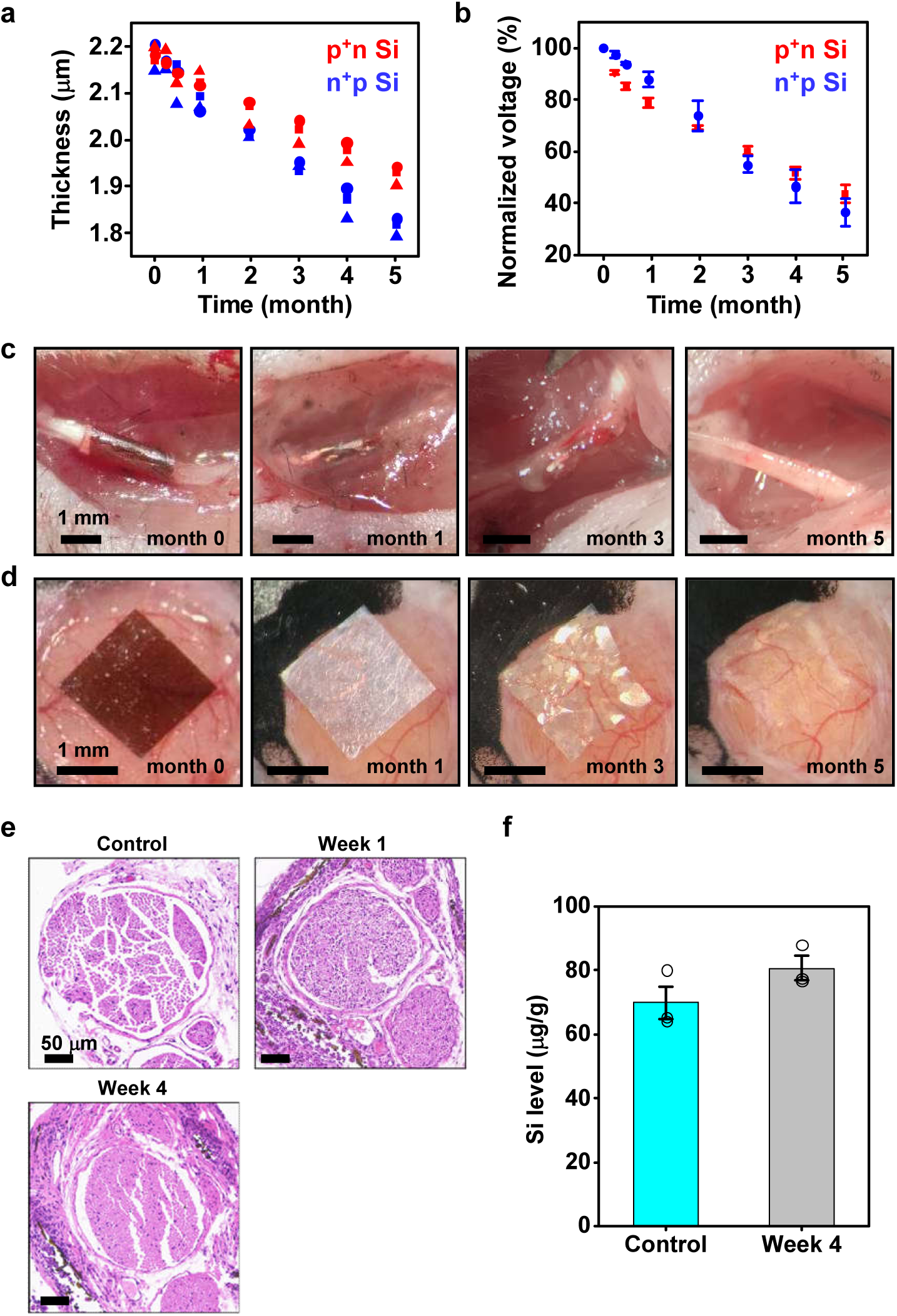
*In vitro* and *in vivo* degradation and biocompatibilities of Si films. (a, b) Changes of (a) thickness and (b) normalized photovoltage for p^+^n and n^+^p Si films immersed in PBS (0.1 M, pH 7.4, 37 °C). Photovoltages are measured under a constant irradiance of 0.9 W/cm^2^. *n* = 3 samples for each film. (c) Series of images showing natural dissolution of a Si film on PLLA-PTMC wrapped around the sciatic nerve of mice. (d) Series of images showing the natural dissolution of a Si film on the brain cortex of mice. (e) Hematoxylin and eosin (H&E) stained histological sections of the sciatic nerve for mice with wrapped Si films implanted for weeks 1 and 4. The image for the control sample is from a mouse without implants. (f) Measured Si concentration in the brain tissue with implanted Si films (*n* = 3 mice). Results for Au are below the detection limit (< 50 ng/g) and not shown here. Control data are from the mice without implant (*n* = 3 mice). Data are presented as means ± s.e.m.

## Discussion

While Si and other materials based devices have been extensively studied for wearable and implantable bio-sensing and modulation^59, 63, 64^, the polarity-dependent optoelectronic response of semiconductor junctions, which have been utilized to produce high-efficiency solar cells and photodetectors for decades, has rarely been exploited for selective neural interrogation. This study demonstrates an unconventional optoelectronic biointerface, simply based on thin-film Si pn diodes, realizing deterministic and non-genetic excitation and inhibition of neural activities in cultured cells, peripheral and central nervous systems. As a practical manner, these Si films can be adapted for wireless, battery-free and lead-free stimulation within the tissue, optically activated by a near-infrared source operated in the tissue transparency window (800–1000 nm). For our flexible Si membranes, readily applied areas include neural modulations in shallow, subcutaneous tissues such as the peripheral nerve^33^, the spinal cord^65^ and the vagus nerve^66^. Another viable application would be the retinal implant^67^. Although optical penetration into the deep tissue is inherently limited to a few millimetres even in the near-infrared window, light delivery can be facilitated by implantable fibers or waveguides, as widely used in optogenetics research.

Our current work employs planar p^+^n and n^+^p Si films, which are suitable for regulating neural cells or tissues over a large area at the millimetre scale. Compared to state-of-the-art optogenetic techniques that can achieve cellular level modulation with high cell specificity^68^, the spatial resolution for these planar Si films is limited by their geometries as well as the light spot size. Nevertheless, one can envision that lithographically patterned and selectively doped n and p regions can be attempted for spatially resolved excitation and inhibition functions in the future^67^. As a preliminary investigation, we numerically study the influence of Si film size on the spatial resolution for optoelectronic stimulation (Fig. S30). It is observed that photogenerated voltages are more focused for films with smaller diameters, which are qualitatively consistent with previous reports^69^. Further reduced device geometries (for example, Si nanorods or nanowires that can be uptaken by cells^70^), combined with a smaller laser beam, can be utilized for more specific cell targeting. Moreover, textured and surface modified Si films (e.g. rough and nanoporous surfaces) can be exploited for enhanced photon responses, reduced junction impedance and improved bio-adhesion^71^. Another possibility is to incorporate such diode designs into self-assembled, 3D scaffolds^72, 73^ for optimal integration with complex biological environments. In addition to the natural dissolution of Si films, encapsulation techniques could also be explored to realize a more controllable degradation process with sustained optoelectronic functions in a predefined timeframe.

When implementing in biological systems, the device platform serves a tool to optically and non-genetically interrogate neural activities, with possible utilities in drug screening^74^, brain-machine interfaces ^75, 76^ and prosthesis^77^. These wirelessly generated electrical signals can also possibly interrogate neurological disorders^78^, guide stem cell migration, modulate progenitor cell development^79, 80^ and promote cell growth and regeneration^81, 82^. Additionally, the junction-dependent photovoltage could target electro-sensitive designer cells expressing voltage-dependent receptors for more cell-specific stimulations^83^. Besides neurons, the optoelectronic stimulation can also regulate behaviors of other cells like glia^37, 70^, cardiomyocytes^84^, oocytes^32^ and HEK cells^29^. In summary, these material and device strategies provide an advanced optoelectronic biointerface with vast opportunities to realize previously inaccessible applications in fundamental biological research as well as clinical medicine.

## Methods

### Preparation of Si pn junctions

Thin-film Si p^+^n and n^+^p diodes were prepared by ion implantation into silicon-on-insulator (SOI) wafers. The p^+^n junction was made of an SOI wafer with an n-Si device layer (100 orientation, resistivity 1–10 Ω*cm, thickness ∼2 µm, Soitec) as the substrate followed by subsequent implantation of boron (B) (dose 4×10^14^ ions/cm^2^, energy 30 keV), while the n^+^p junction was made of an SOI wafer with a p-Si device layer (100 orientation, resistivity 1–10 Ω*cm, thickness ∼2 µm, Soitec) as the substrate followed by subsequent implantation of phosphorus (P) (dose 4×10^14^ ions/cm^2^, energy 75 keV). After cleaning, these implanted wafers were annealed for 30 mins at 950 °C for dopant activation.

### Fabrication of Si films on flexible substrates

Patterns of doped Si films were lithographically defined by reactive ion etching (power 100 W, 150 sccm SiF6, 90 mTorr pressure, etch rate 20 nm/s). The samples were cleaned in H_2_O:H_2_O_2_:NH_4_OH = 5:1:1 (10 mins, 80 °C), and hydrofluoric acid (49% HF, ACS grade, Aladdin) removed the buried oxide layer and released the 2 µm thick Si films. Polydimethylsiloxane (PDMS, Dow Corning Sylgard 184 kit, 1:10 weight ratio) stamps and thermal release tapes (No.3198, Semiconductor Equipment Corp.) were applied to transfer the freestanding Si films onto target substrates. As an adhesive layer, an epoxy (SU8-3005, 5 µm thick) was spin coated on carrier substrates before the transfer printing. After transferring Si films, samples were baked at 110 °C for 30 mins and cleaned with acetone, isopropanol and deionized water.

For optical modulations in mice’s sciatic nerve and cerebral cortex (Fig. 4 and 5), Si films were transferred on a highly transparent polyethylene terephthalate (PET, thickness ∼25 µm) film, and decorated with gold nanoparticles, by dipping into 0.5 mM HAuCl4 (No.G4022, Sigma-Aldrich) in 1% HF (GR, 40%, Aladdin) for 3 mins. For in vivo degradation tests (Fig. 6), Si films were mounted on a self-adhesive, water-soluble film made of poly(L-lactic acid) and poly(trimethylene carbonate) (PLLA-PTMC^82^) copolymer (thickness ∼100 µm).

### Optoelectronic properties of Si films

A standard patch-clamp setup was used to measure the photon response of the Si films. A red laser (635 nm, spot size ∼2 mm) was incident on the material through a collimating lens, and its irradiance was controlled by transistor-transistor logic (TTL) signals. Voltage-and Current-clamp protocols were performed with an Axopatch 200B amplifier controlled by pClamp software (Molecular Devices). Glass pipettes (∼ 1 MΩ) filled with 1×PBS solution approached to the Si surface with a distance of ∼5 µm. For steady-state photoresponse measurements, a PBS droplet is placed on Si films, of which the backside connects to the ground of the amplifier. Specifically, metal films (Au for p-Si and Al for n-Si) were sputtered on the backside of Si and bonded to a copper sheet with silver paste. The photovoltages were taken by current-clamp recording (filtered at 3.2 kHz and sampled at 10 kHz). The transient photoresponse measurements were taken by voltage- or current-clamp recording (filtered at 10 kHz and sampled at 200 kHz), with Si films fully immersed in PBS solution.

### Simulations for spatial distribution of photovoltages

The spatial distribution of photovoltage generated by Si films within the solution was calculated in a 2D geometry via finite element analysis (COMSOL Multiphysics 5.5), with the Electrical Static (ES) model of the AC/DC module. Si surfaces exposed to solution were assumed to be equipotential surfaces and the values of electrical potential were set to 50 mV and −60 mV for n^+^p and p^+^n Si diodes, respectively. The PBS solution was set as a dielectric with a relative permittivity of 80. The distance from the boundary of electrolyte to the centre of silicon devices was 2 cm and the boundary was fixed at 0 V.

### Photothermal characterizations

Photoinduced heating effects on Si surfaces were measured with a thermocouple (IT-24P, BAT-10R, Physitemp Instruments, LLC). The thermocouple positioned on the Si surface immersed in PBS. The temperature increases were recorded as a function of light intensity (0–5 W/cm^2^) and pulse duration (1–10 s).

### In vitro hydrolysis of Si films

Samples of Si diodes with etched patterns (2 × 2 mm^2^) were immersed into PBS solution (0.1 M, pH 7.4) at 37 °C for up to 160 days. Changes of film thickness were monitored by a profilometer (Alpha-Step D500), and transient photovoltages were taken by current-clamp recording under illumination (635 nm, 0.9 W/cm^2^, 10 ms duration).

### Animal care

All animal protocols used were in accordance with the institutional guidelines of the National Institute of Biological Sciences in Beijing (NIBS), Tsinghua University and the Shenzhen Institute of Advanced Technology (SIAT), and were proved by Institutional Animal Care and Use Committee (IACUC). All animals are socially housed in a 12 h/12 h (lights on at 8 am) light/dark cycle, with food and water ad libitum.

### Culture of DRG neurons

The adult rat DRGs (3−4 months) were extracted and transferred into a small tube containing ice-cold 1 ml of Dulbecoo’s modified eagle medium (DMEM/F12) supplemented with 1% penicillin-streptomycin (PS) solution. Tissues were cut into small pieces and treated with an enzyme solution containing 5 mg/ml dispase and 1 mg/ml collagenase at 37 °C for 1 hr. After trituration and centrifugation, cells were washed in 15% (w/v) bovine serum albumin (BSA) resuspended in DMEM/F12 containing 10% fetal bovine serum (FBS), and 1% PS dropped on Si and glass materials coated with 0.01% poly-L-lysine (PLL, No.P8920, Sigma-Aldrich) and 1–2 μg/cm^2^ laminin (No.L2020, Sigma-Aldrich), cultured in an incubator at 37 °C, and used after 2 days.

### In vitro electrophysiological recording

DRG neurons were perfused with Hanks balanced salt solution (HBSS) contained the following (in mM): 137 NaCl, 0.6 EGTA, 5.4 KCl, 0.4 MgSO_4_, 0.5 MgCl_2_, 1.3 CaCl_2_, 5.5 Glucose, 4.2 NaHCO_3_ and 0.44 KH_2_PO_4_ (pH 7.2–7.4) at a rate of 3 ml/min at room temperature. Cells were identified with differential interference contrast optics (DIC; Zeiss Examiner.Z1). Recording pipettes (3–6 MΩ) were pulled with a micropipette puller (P1000, Sutter Instrument; USA). For whole-cell recordings, pipettes were filled with internal solution that contained the following (in mM): 130 K-gluconate, 10 HEPES, 0.6 EGTA, 5 KCl, 3 Na_2_ATP, 0.3 Na_3_GTP, 4 MgCl_2_, and 10 Na_2_-phosphocreatine (pH 7.2–7.4). Voltage- and current-clamp recordings were performed with a computer-controlled amplifier (MultiClamp 700B, Molecular Devices; USA). Recorded traces were low-pass filtered at 3 kHz and sampled at 10 kHz (DigiData 1440, Molecular Devices). Collected data were analyzed using Clampfit 10 software (Molecular Devices). Light stimulations were generated by a 635 nm laser (Thorlabs), with controlled power, frequency and duration (Digidata 1440, Molecular Devices).

### Circuit model for cell stimulation

A circuit model (simulated by Multisim 14.1) was established to understand the mechanism of extracellular photocapacitive stimulation^49^. The cell was assumed to have a hemispheric shape with a diameter of 30 µm. Parameters of circuit elements such as membrane resistance and capacitance were adopted from references^85,86,87^. Photoinduced inward or outward transmembrane currents were taken and normalized based on experimental data (Figs. S12 and S13).

### Calcium imaging

Cells were loaded with 10 mg/ml Fluo-4 AM (Thermo Fisher Scientific) supplemented with 0.01% Pluronic F-127 (w/v; Invitrogen), at 37 °C for 1 hr. Subsequently, cells were continuously perfused in HBSS at a rate of 3 ml/min at room temperature (25 ± 2 °C). Ca^2+^ fluorescent signals were imaged with a 20× water immersion objective on a confocal microscope (FV1000, Olympus). A 635 nm laser was incident on the samples and controlled by a programmable stimulator (Master 4, A.M.P.I.). For experiments of cell inhibition, 4 μM α-amino-3-hydroxy-5-methyl-4-isoxazolepropionic acid (AMPA, No.A6816, Sigma-Aldrich) was initially added in HBSS buffer to enhance baseline Ca^2+^ signals. Fluorescence images were analyzed via ImageJ. Normalized fluorescence changes were calculated as Δ*F*/*F* = (*F* − *F*_0_) / *F*_0_, where *F*_0_ is the baseline intensity (before 5 s).

### Scanning electron microscopy

DRG neurons cultured on Si and glass were fixed in mixture of 2% paraformaldehyde (PFA) + 2.5% glutaraldehyde (GA) solution, and then gradiently dehydrated with ethanol and dried with tert-butyl alcohol (TBA). A thin layer of gold was sputtered on samples, and images were taken with FEI Quanta 200 SEM.

### Stimulating and recording activities of the sciatic nerve in vivo

Wild-type mice (C57BL/6, 6 weeks old) were purchased from VitalRiver (Beijing, China). Animals were anesthetized with 2% isoflurane in balanced oxygen during the operative procedure. The sciatic nerve was exposed without muscle damage by releasing surrounding connective tissue. Si films were transferred onto PET substrates and attached on the fiber surface. Through a collimating lens, a red laser beam (635 nm, ∼2 mm spot size) illuminated the Si film, controlled by transistor-transistor logic (TTL) signals. A multi-channel electrophysiological acquisition and processing system (Chengdu Instrument Factory, RM6240E/EC) recorded and stimulated muscular activities. A recording electrode was inserted into the tibial related-muscle region, collecting electromyography (EMG) signals with a sampling rate of 20 kHz and a low-pass filter at 300 Hz. During inhibition experiments, another stimulating electrode positioned at the proximal site of nerve evoked CMAPs, with a frequency of 4 Hz, a pulse voltage of 4 mV and a pulse width of 1 ms. A camera captured the hindlimb lifting with a sampling rate of 30 frames/s. Deeplabcut (resnet50) was used to obtain the coordinates of the farthest position of the mouse leg in each frame.

### Stimulation and recording activities of the brain cortex in vivo

Wild-type mice (C57BL/6, 2−3 months) were deeply anaesthetized with pentobarbital sodium (80-100 mg/kg) and placed in a stereotaxic frame. A skin incision was performed over the cerebral cortex to expose the skull. A stainless-steel set screw, together with a spade terminal was affixed to the skull with dental cement, hardened for at least 30 mins. Subsequently, the screw was mounted into the optical post to fix the mouse head for surgical operations. A dental drill created a hole with a diameter of ∼3 mm over the motor and somatosensory cortices and then the dura was peeled off to fully expose the cortex. During experiment, 0.9% saline was frequently applied to the exposed cortex area to prevent dehydration. Before recordings, animals were transitioned from pentobarbital sodium to isoflurane to reach a more stable state of anaesthesia. A customized laser scanner (473 nm, ∼2 mm spot size) delivered light on Si films attached on the cortex. Electrophysiological signals were recorded by a 32-channel silicon probe (Lotus). The implanted probe was tilted by 40° off the vertical axis to collect neural signals underneath the Si film. The depth of probe tip was ∼800 μm from the pia mater, with the entry point in the somatosensory cortex close to the edge of the Si film. Data were collected at 20 kHz, with a set of bandpass filters (Butterworth, third order, 300–6000 Hz) and a 50 Hz notch filter in each neural channel. A PCA filter was applied to minimize photoinduced artifacts, and the threshold was set to four times the standard deviation to detect spikes. Data were analyzed with customized Matlab codes.

### In vivo degradation of Si films

Si films with a dimension of 2 mm × 2 mm × 2 µm were implanted into wild-type mice (C57BL/6, 6 weeks old, purchased from VitalRiver, Beijing, China). During the operation, animals were anesthetized with 2% isoflurane in balanced oxygen. For implants around the sciatic nerve, the nerve fiber was exposed without muscle damage by releasing surrounding connective tissue. The Si film was integrated onto a self-adhesive biodegradable copolymer of poly(L-lactic acid) and poly(trimethylene carbonate) (PLLA-PTMC) film, wrapped around the sciatic nerve, and fixed with 8-0 silk suture. The skin incision was closed with 5-0 silk suture. For implants in the brain, the Si film was placed on the cortex after opening the skull. A glass window replaced the removed skull for tissue protection and clear optical imaging. The degradation of Si films on the sciatic nerve and the cortex was monitored by a camera at a regular period of time. For histological imaging, sciatic nerves of mice were extracted (∼3 mm length) and fixed with 4% paraformaldehyde in PBS overnight. Subsequently, standard methods of embedding, slicing and hematoxylin-eosin (H&E) staining were imposed. For spectroscopic characterization of the elemental distributions (Si and Au) in living tissues, brains of mice were extracted after perfusion with PBS. The brain tissues were fully dried by vacuum freeze dryer (ALPHA, Christ) for about 6 hours after pre-freeze step at −80 °C overnight. After weighting, the samples were transferred to a clean microwave digestion tank and added with nitrohydrochloric acid (nitric acid: hydrochloric acid = 3:1) and placed overnight. The samples were microwave digested (160 °C, 25min). After cooling, the dissolved tissue solutions were analysed by ICP-OES (IRIS Intrepid II, Thermo) for Si element and by ICP-MS (XSeries II, Thermo) for Au element, respectively. The blood (∼200 μL per mouse) was collected into K_2_-EDTA container by capillary tube from the orbital vein of the isoflurane anesthetized mice. Hematology analyses were conducted by Charles River Laboratories.

## Data availability

The main data supporting the results in this study are available within the paper and its Supplementary Information. The raw data acquired during the study are available from the corresponding author on reasonable request.

## Code availability

The Matlab and Deeplabcut codes used in this study are available from the corresponding author on request.

## Acknowledgements

This work is supported by National Natural Science Foundation of China (NSFC) (61874064), Beijing Municipal Natural Science Foundation (4202032), Beijing Innovation Center for Future Chips at Tsinghua University, Center for Flexible Electronics Technology at Tsinghua University, Beijing National Research Center for Information Science and Technology (BNR2019ZS01005), and National Key R&D Program of China (2018YFA0701400).

## Author contributions

Y. H. and X. S. developed the concepts. Y. H., H. W., Y. X., P. S. and X. F. performed material design, fabrication and characterization. Y. H., H. W., J. C., L. L. and X. S. performed numerical simulations. Y. H., Y. C., H. D., J. W., R. H., S. H., H. H., D. Y., X. F., S. W. and X. L. designed and performed biological experiments. L. Y., W. X., M. L., S. S., S. W., X. L. and X. S. provided tools and supervised the research. Y. H. and X. S. wrote the paper in consultation with other authors.

---

**Figure S1.**
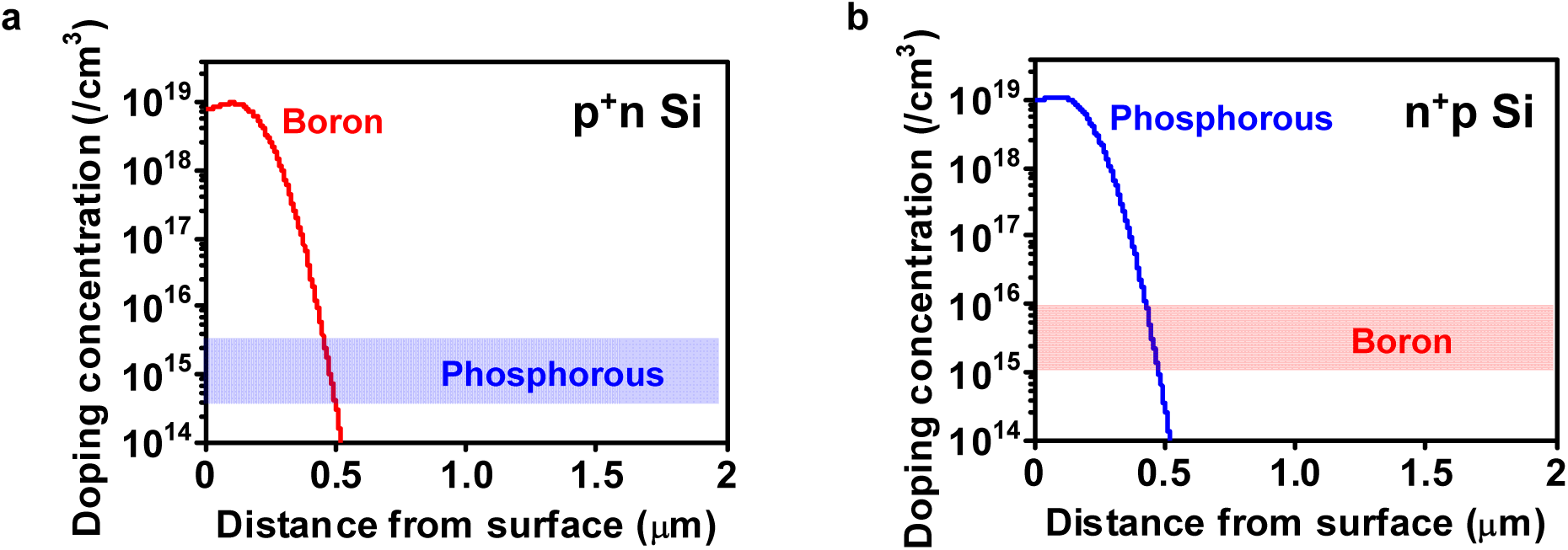
Designed doping profiles for Si films made of (a) p^+^n and (b) n^+^p junctions.

**Figure S2.**
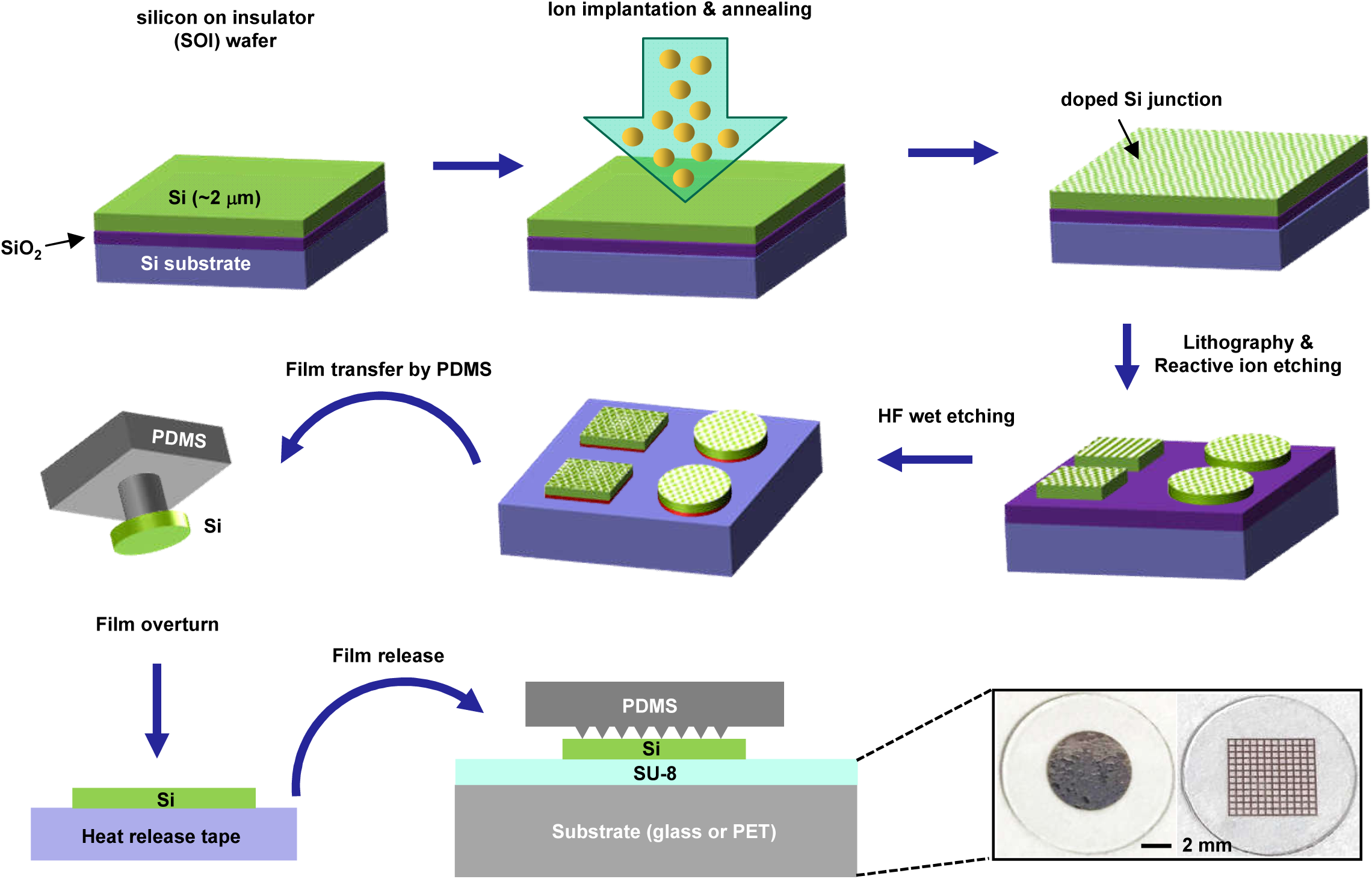
Schematic illustration of processing flow for the fabrication and transfer printing of doped silicon membranes.

**Figure S3.**
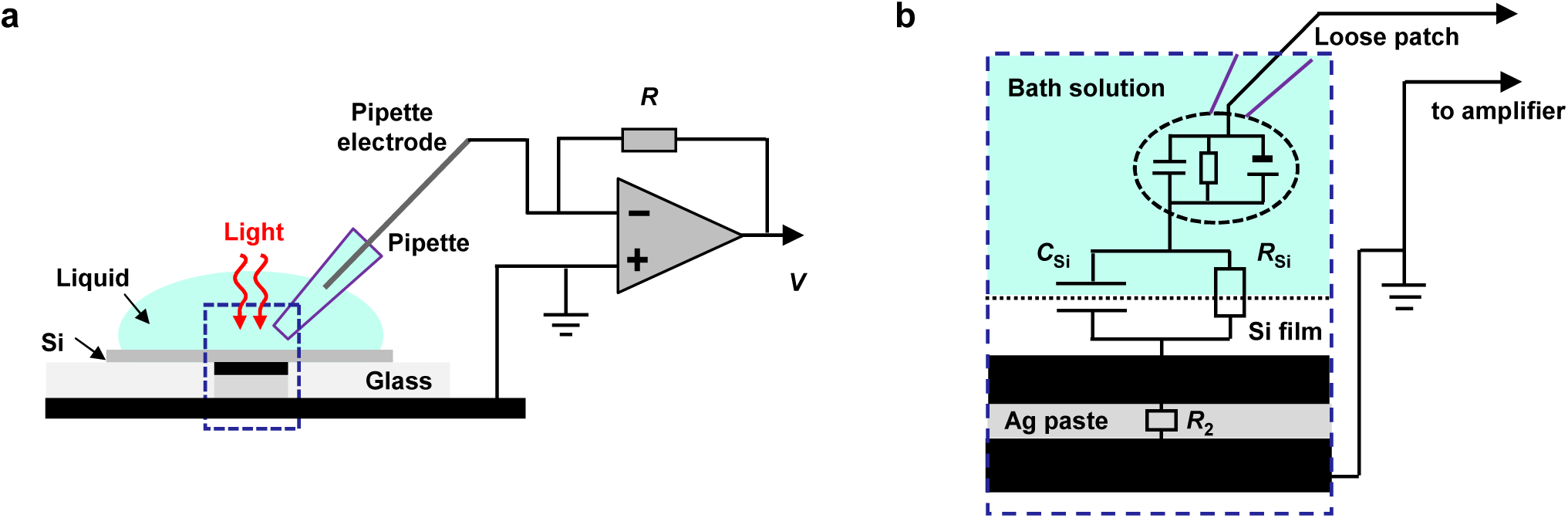
(a) Detailed schematics for the setup of steady-state photoresponse measurement. (b) Corresponding circuit diagram. Photovoltages are taken by current-clamp recording (filter at 3.2 kHz and sampled at 10 kHz) and the resistance of pipette is ∼1 MΩ. The contact electrode of the back belonged to the p-type Si is sputtered with Au and the n-type Si is sputtered with Al. The copper sheet wired to the circuit of the amplifier attaches to the contact electrode with Ag paste.

**Figure S4.**
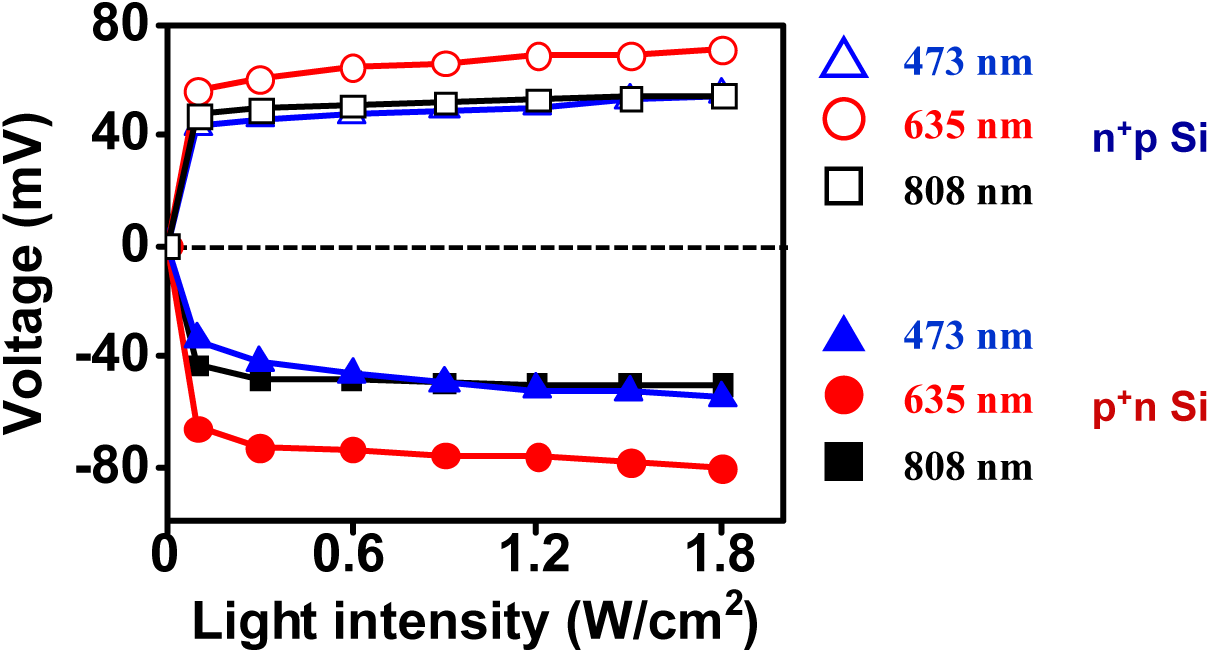
Measured steady-state photovoltages versus light intensity under illumiation at different wavelengths (473 nm, 635 nm, and 808 nm) with the n^+^p and p^+^n Si films during 5 s continuous illumination.

**Figure S5.**
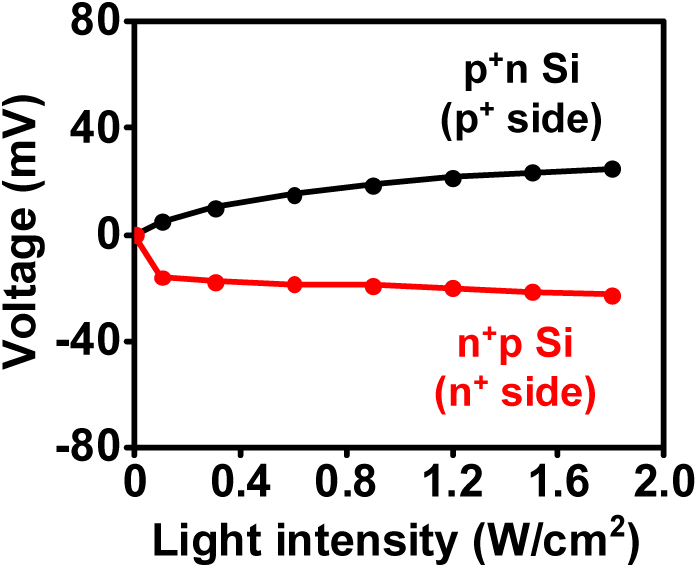
Measured steady-state photovoltages versus light intensity n^+^p and p^+^n Si films, with the highly doped regions facing up and contacting with solutions.

**Figure S6.**
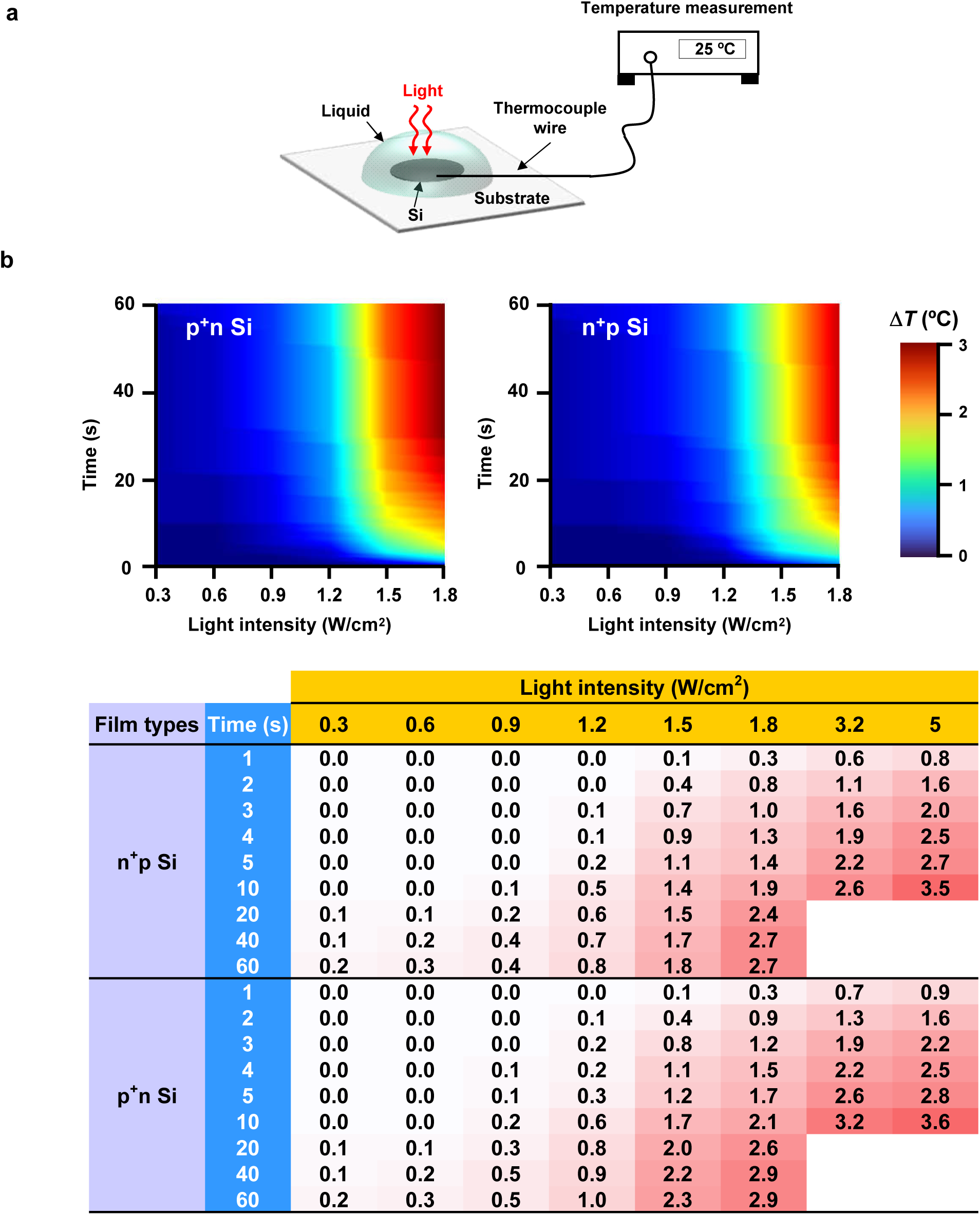
(a) Setup of temperature measurements for Si films in solutions under light illumination. (b) Measured maximum temperature rises (unit: °C) on the Si surface, as a function of the illumination duration time and the light intensity.

**Figure S7.**
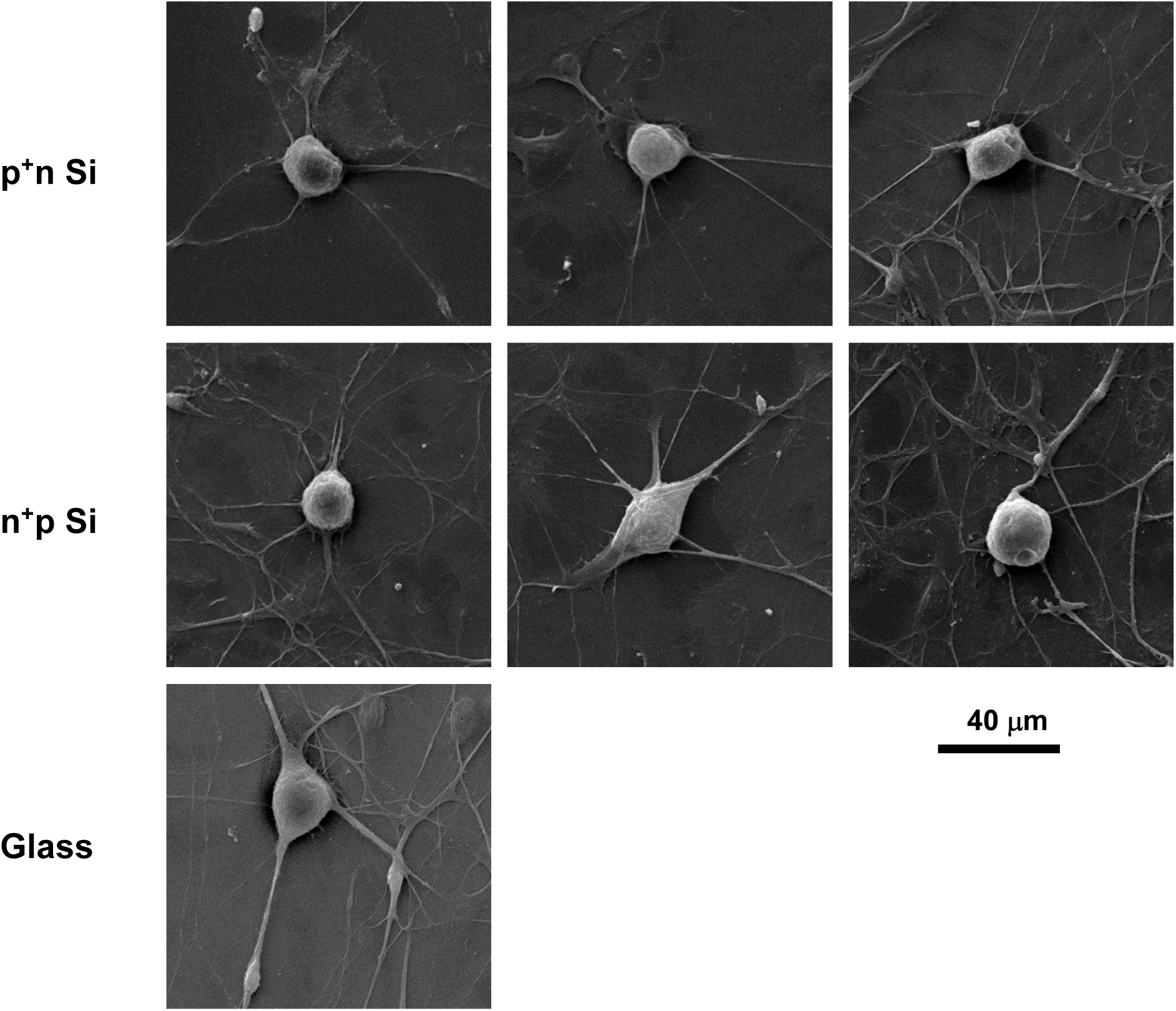
SEM images of DRG neurons cultured on Si films and glass, indicating the large size DRG neurons (20–40 μm) and the small-size glial cells (5–10 μm).

**Figure S8.**
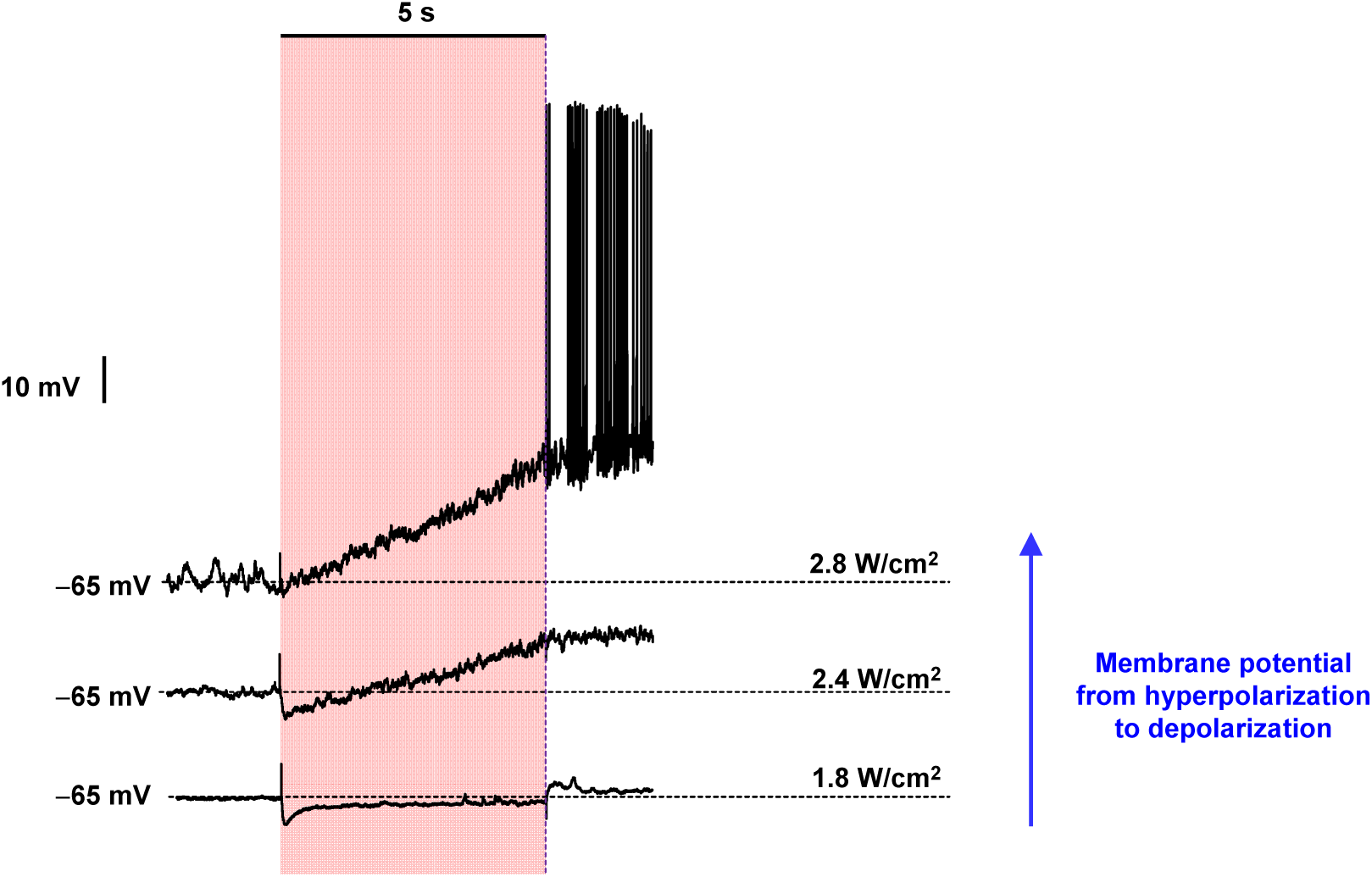
Transition from cell hyperpolarization to depolarization with the n^+^p Si film, by continuous illumination with increased intensities. The spikes appear immediately after the illumination at 2.8 W/cm^2^ for 5 s.

**Figure S9.**
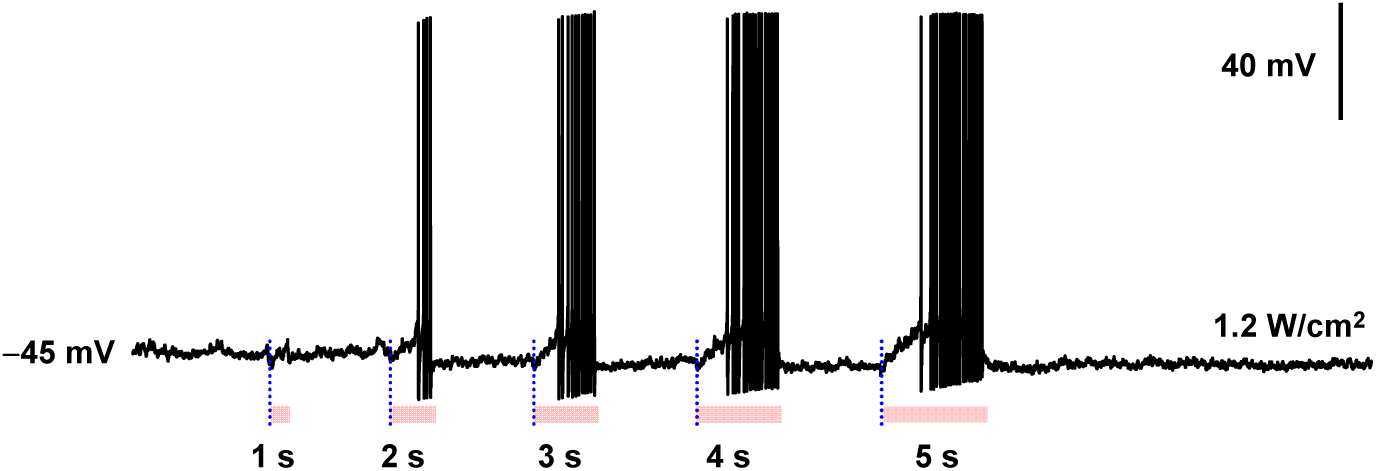
Typical traces from current-clamp recordings of the neural action potentials caused by depolarization effects of the p^+^n Si with various stimulating durations at 1.2 W/cm^2^.

**Figure S10.**
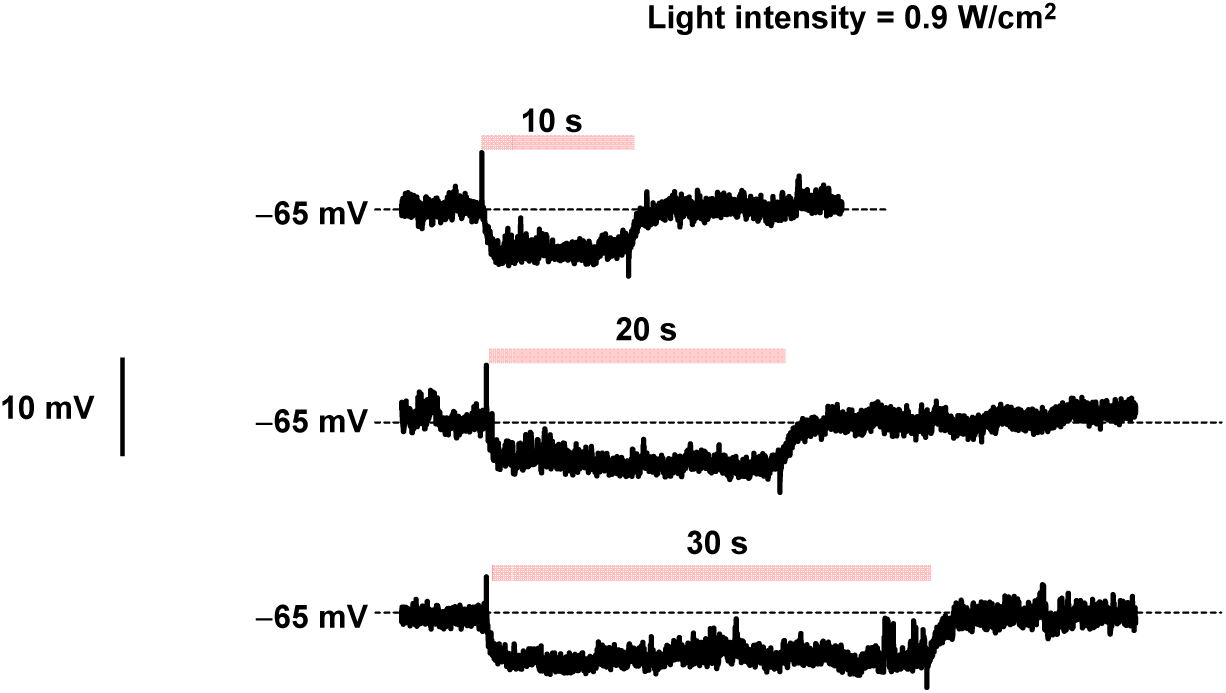
Hyperpolarization effects of the n^+^p Si by continuous illumination at an intensity of 0.9 W/cm^2^ for 10 s, 20 s and 30 s.

**Figure S11.**
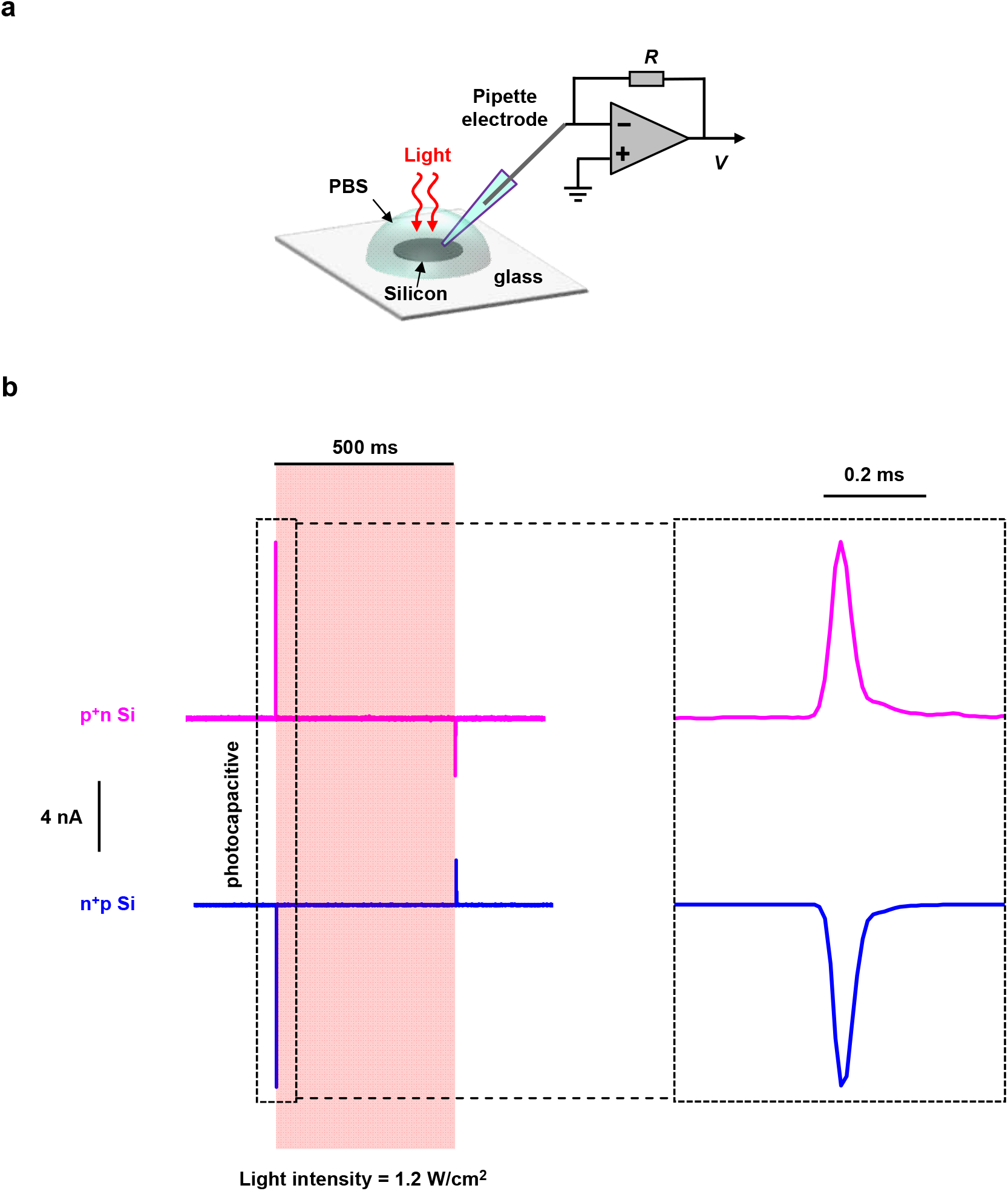
Measurement of photocurrents induced by p^+^n and n^+^p Si diode films. (a) Scheme of the setup for the transient photoresponse measurement. The transient photocurrents are taken by voltage-clamp recording (filter at 10 kHz and sampled at 200 kHz) and the resistance of pipette is ∼1 MΩ. Lightly doped surfaces (n-side for p^+^n Si, and p-side for n^+^p Si) contact the solution. (b) The transient photocurrents generated on the Si surfaces, with pulsed light duration 500 ms and intensity 1.2 W/cm^2^. Most of the currents are photocapacitive, and negligible Faradic currents are observed.

**Figure S12.**
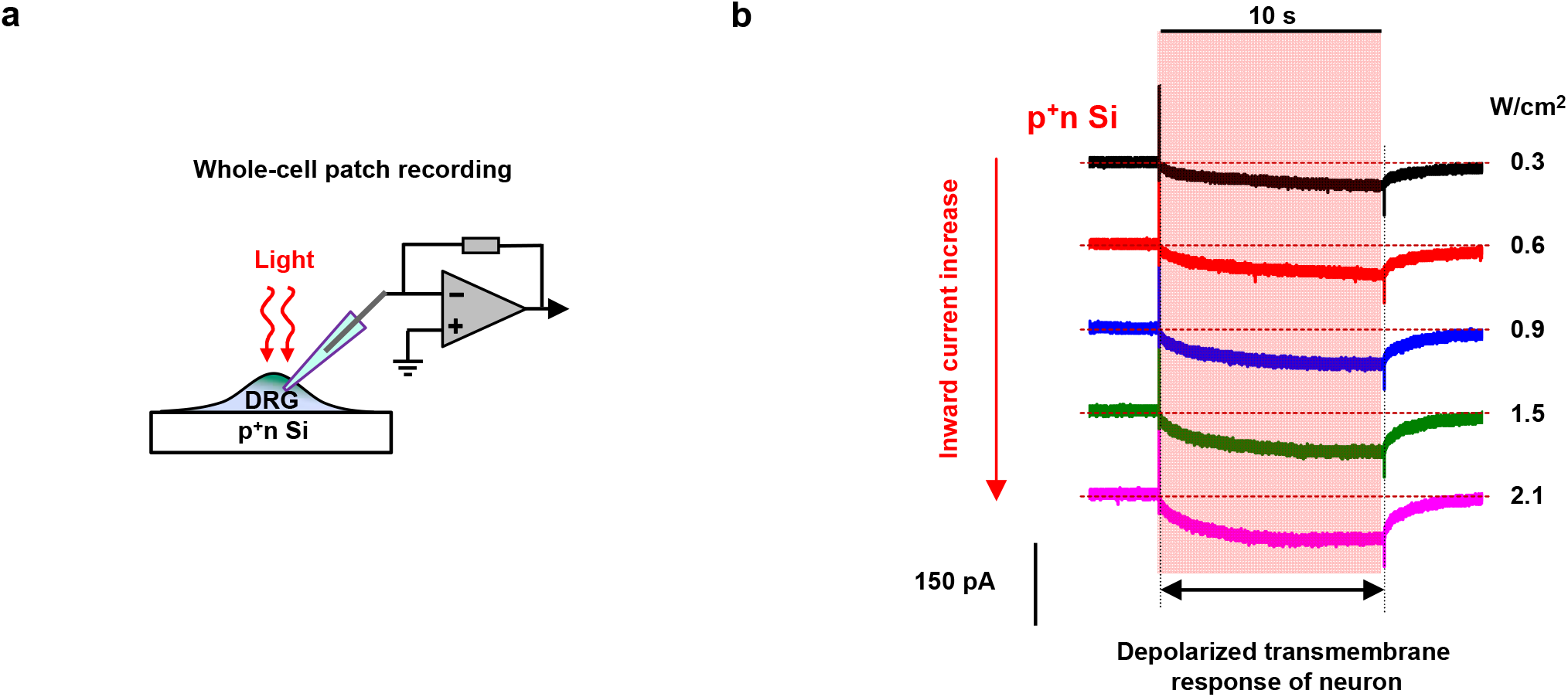
Depolarized membrane current with the p^+^n Si film. (a) Scheme of the setup for the whole-cell recording of the DRG grown on p^+^n Si. (b) Typical recorded membrane currents in response to the photostimulation with varying light intensities within a 10-s-long illumination, showing the slowly increased depolarized currents.

**Figure S13.**
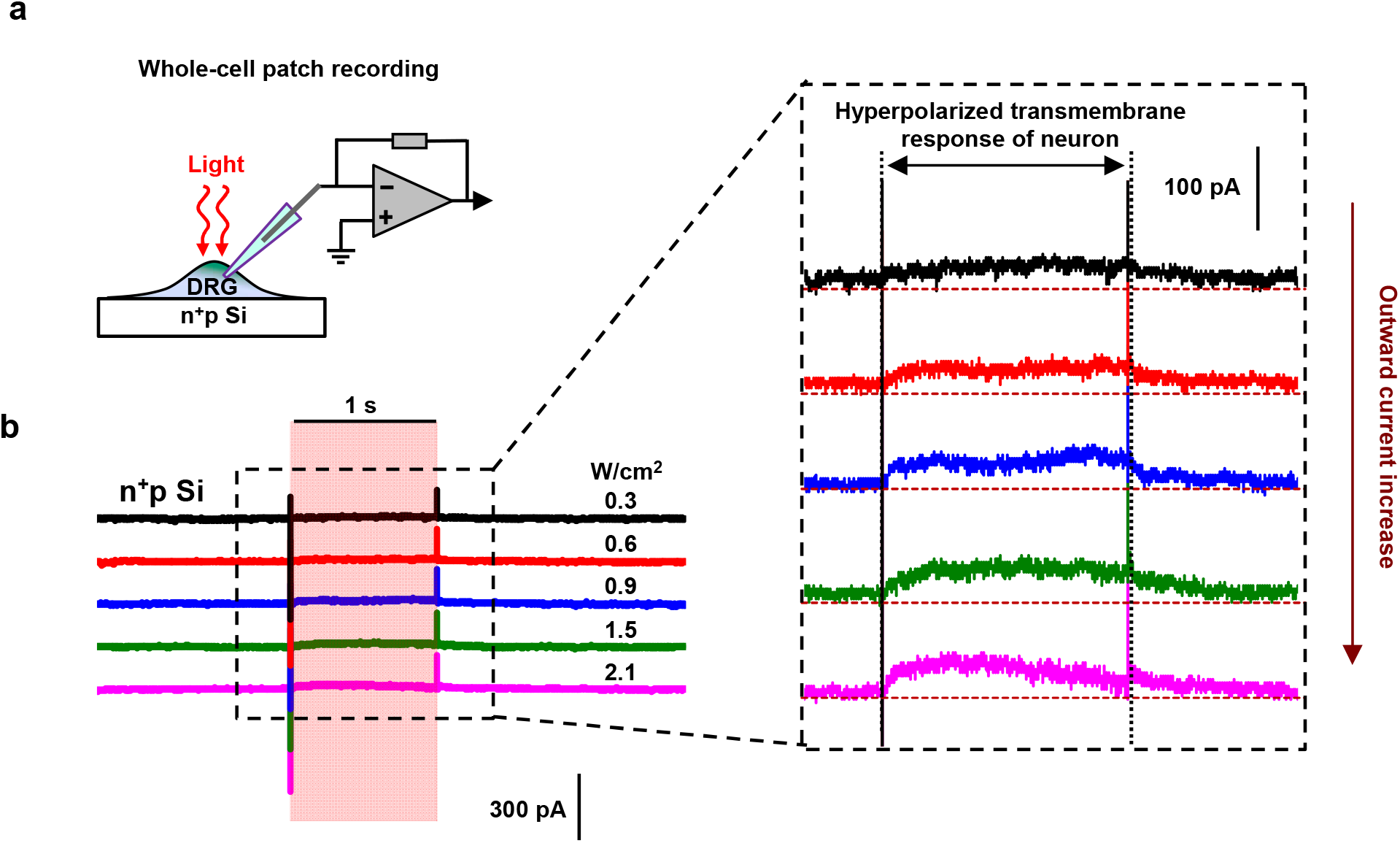
Hyperpolarized membrane current with the n^+^p Si film. (a) Scheme of the setup for the whole-cell recording of the DRG grown on n^+^p Si. (b) Typical recorded membrane currents in response to the photostimulation with varying light intensities within a 1-s-long illumination (left), and enlarged details of the hyperpolarized currents (right).

**Figure S14.**
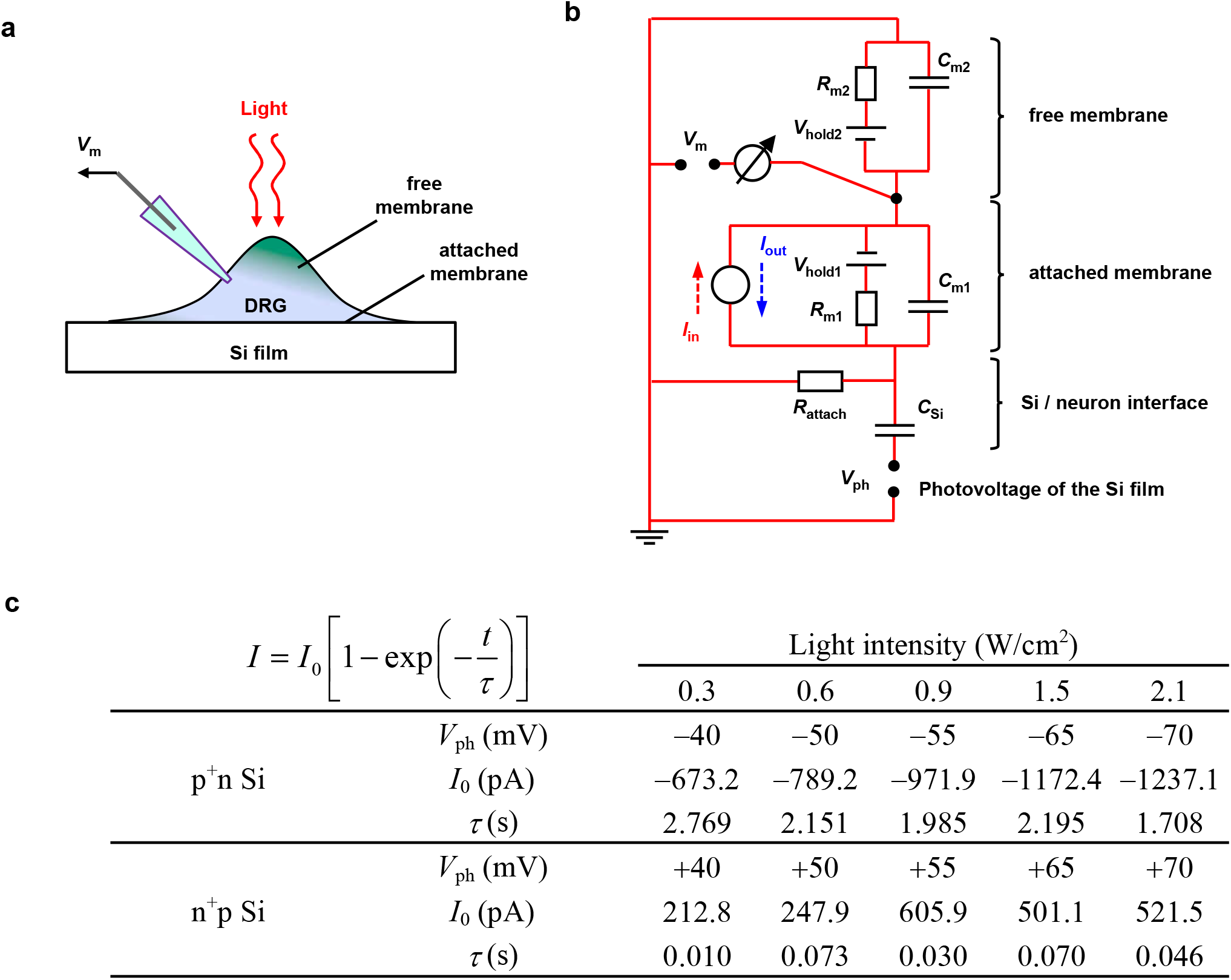
Circuit model developed to understand the polarity-dependent photostimulation. (a) Scheme of the setup for the whole-cell patch clamp recording of DRG cells cultured on Si films. (b) Equivalent circuit of the recorded membrane potential influenced by photovoltaic stimulations. Here we assume the cell has a hemispheric shape with a diameter of 30 µm. *R*_m1_ = 66.9 MΩ, *R*_m2_ = 33.45 MΩ, *C*_m1_ = 27.3 pF, *C*_m2_ = 54.6 pF, *C*_Si_ = 70.7 pF and *R*_attach_ = 1.4 kΩ. The initial holding potential *V*_hold_ = −65 mV. (c) Parameters used for photovoltages *V*_ph_ generated by p^+^n and n^+^p Si films (taken from Figs. 1b and 1c), as well as transmembrane inward and outward currents *I*_in_ and *I*_out_ (taken and normalized from Figs. S12 and S13), in response to various light intensities.

**Figure S15.**
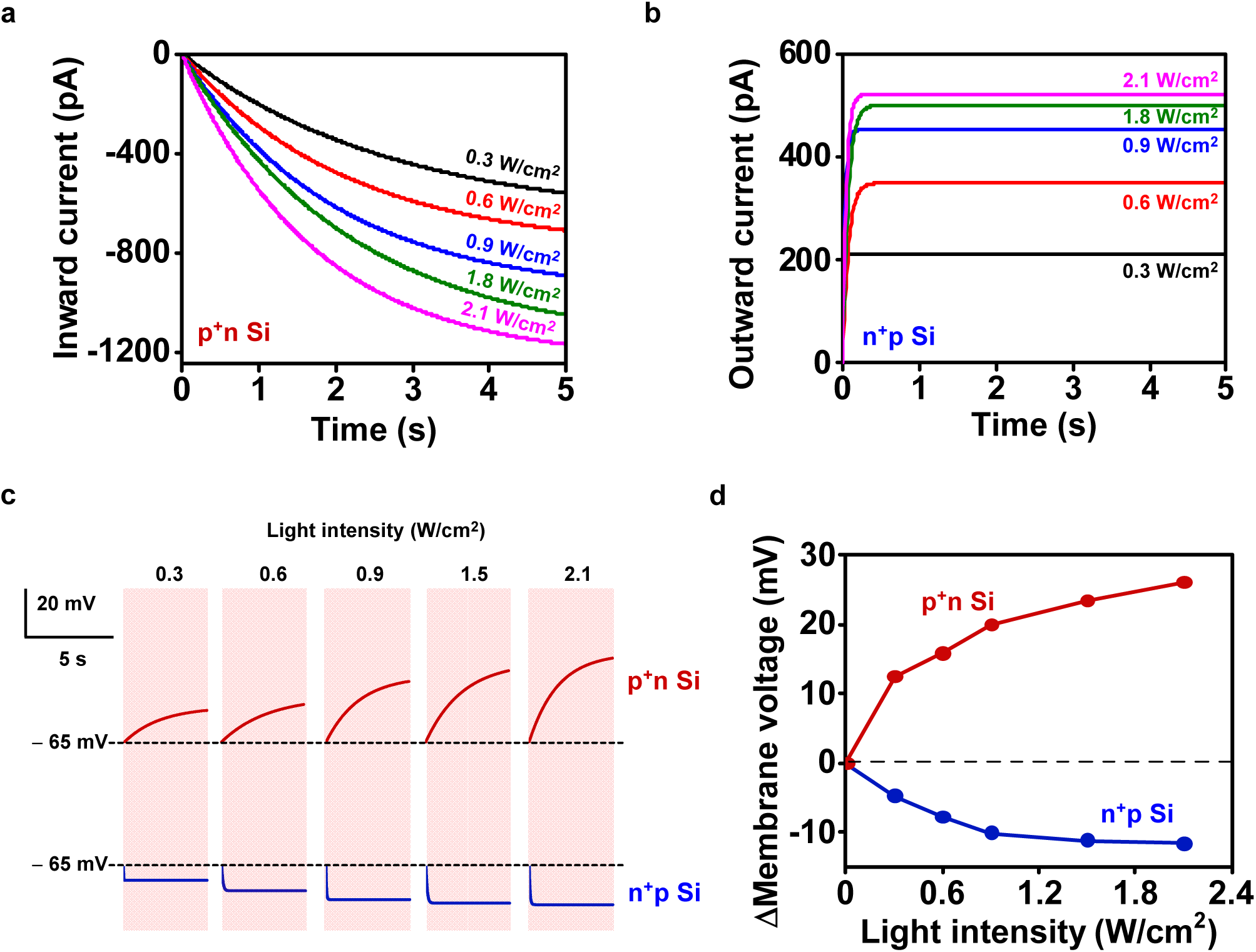
Simulation results based on the circuit model. (a) Inward and (b) outward transmembrane currents (*I*_in_ and *I*_out_) applied in the model, based on parameters in Fig. S14. (c) Calculated membrane voltages (*V*_m_) responding to different inward and outward currents generated by p^+^n Si and n^+^p Si, respectively. (d) Depolarized and hyperpolarized membrane voltages as a function of the light intensity. The simulation results are in good agreement with experiments in Figs. 2c and 2d.

**Figure S16.**
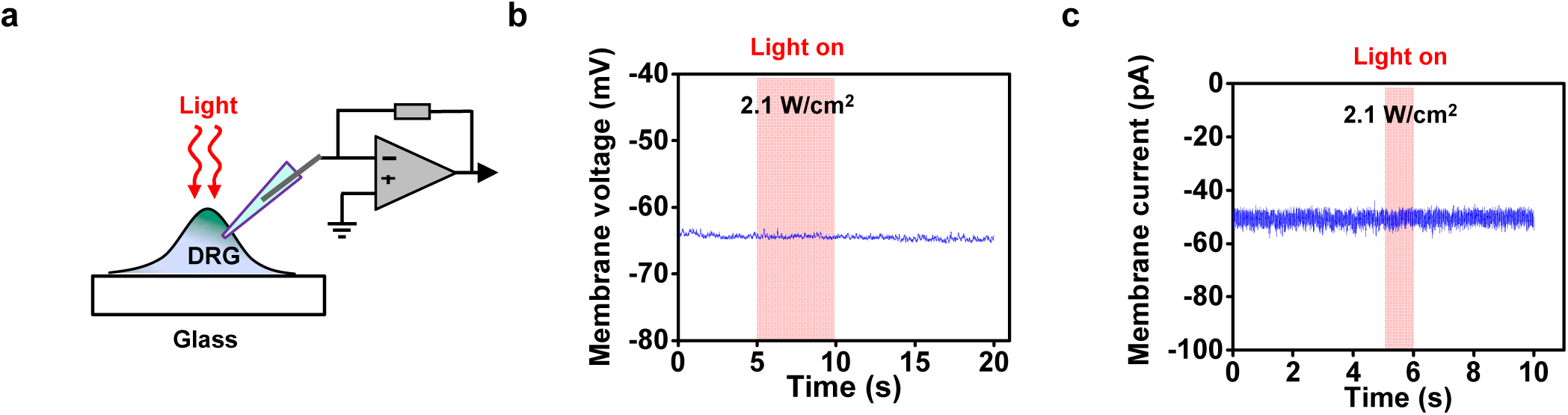
Photostimulated neurons on the glass substrate. (a) Scheme of the setup for the photoresponse measurement. (b) Example trace of the recorded membrane voltage with holding initially at −65 mV and during 5-s-long light stimulation. (c) Example trace of the recorded membrane current with initial potential at −65 mV and during 1-s-long light stimulation. More than 5 independent neurons are taken and each one is repeated 3 times.

**Figure S17.**
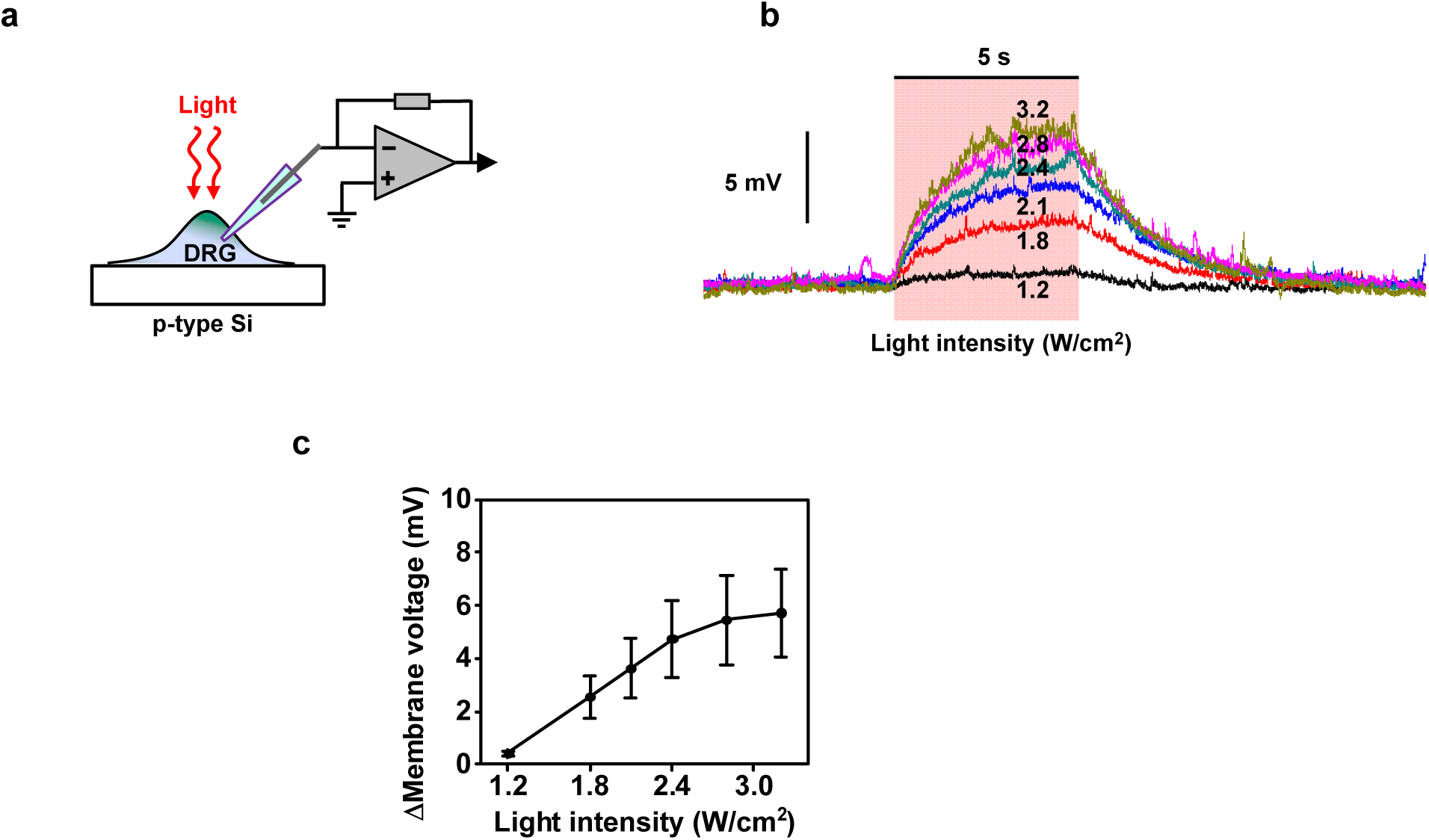
Photostimulated neurons based on the p-Si film. (a) Scheme of the setup for the photoresponse measurement. (b) The typical membrane voltage responses with different light intensities. (c) Statistics of the changed membrane voltages with different light intensities during a 5-s-long illumination (*n* = 5 neurons). These small photogenerated depolarizations are due to the band bending by p-type Si semiconductor-solution junction. All data are presented as means ± s.e.m.

**Figure S18.**
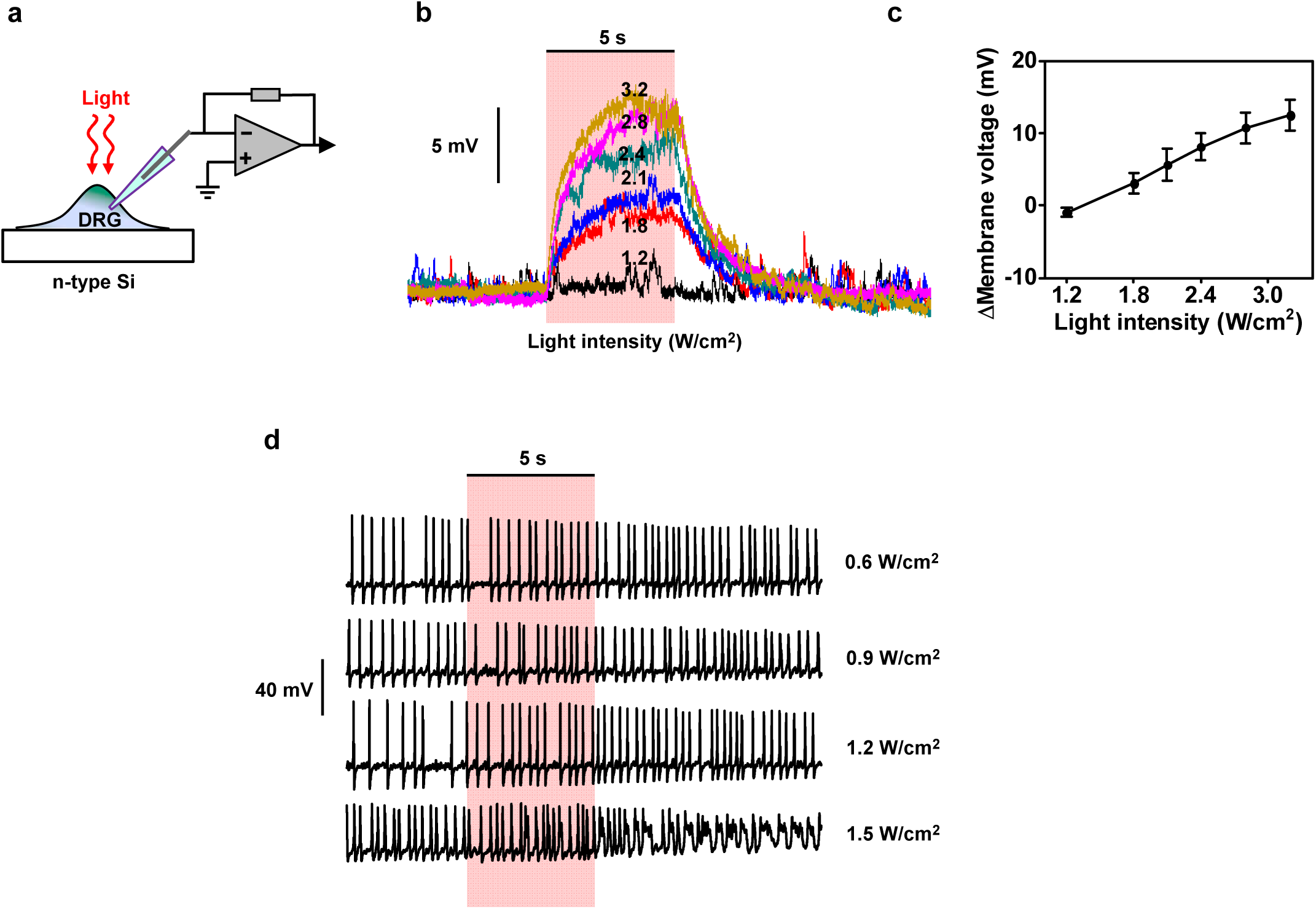
Photostimulated neurons based on the n-Si film. (a) Scheme of the setup for the photoresponse measurement. (b) The typical membrane voltage responses with different light intensities. (c) Statistics of the changed membrane voltages with different light intensities (*n* = 5 neurons). These photogenerated depolarizations are due to the band bending by n-type Si semiconductor-solution junction. Data are presented as means ± s.e.m.. (d) Example traces of the silent effects for an excited neuron at different light intensities (*n* = 2 neurons). No obvious neuron inhibition effects are observed.

**Figure S19.**
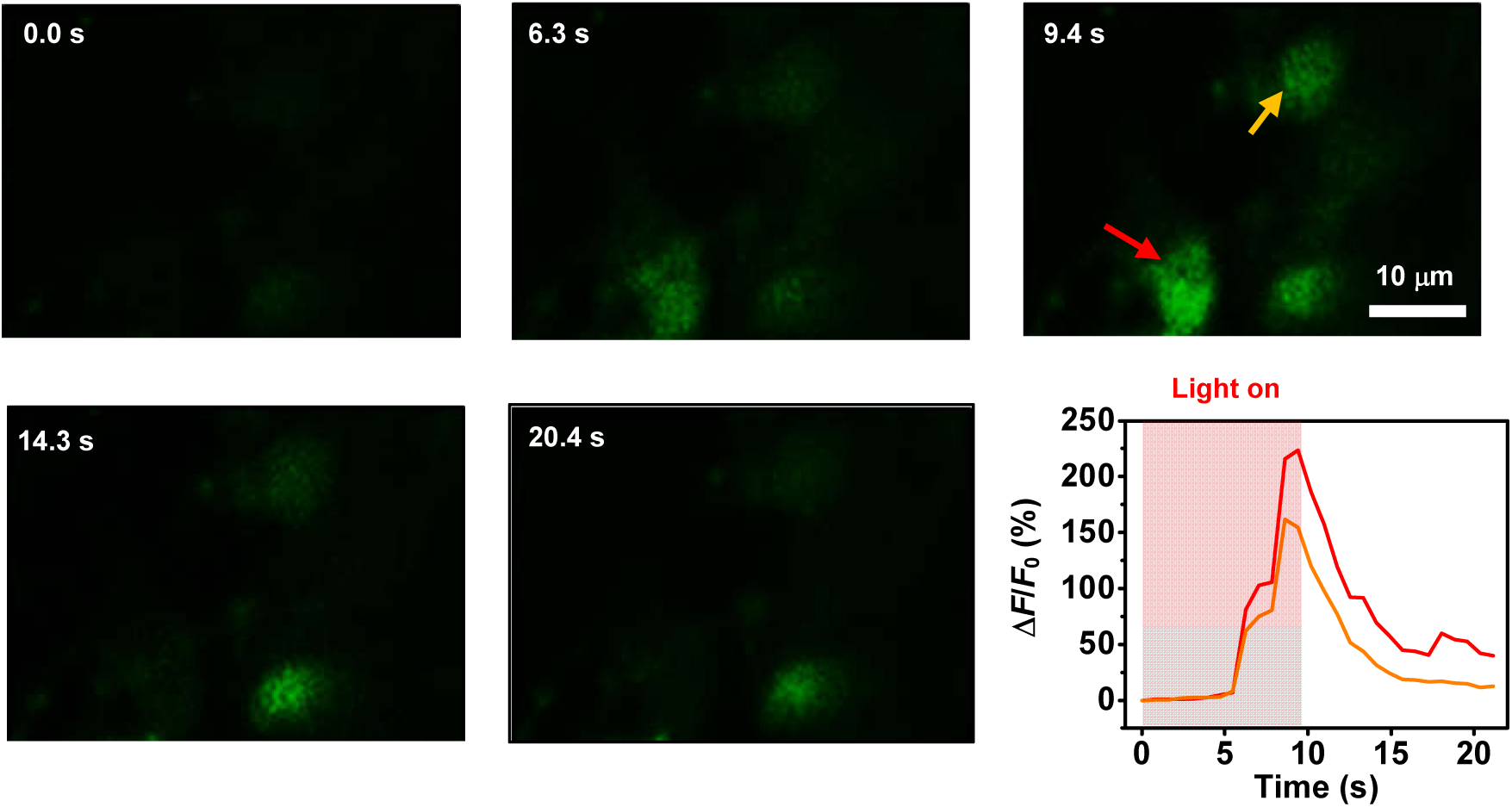
Calcium imaging of glial cells on p^+^n Si films. Glial cells can be optically activated to trigger intracellular calcium elevation (green) over time. The arrow mark the glial cells. The light is on from 0 s to 10 s (630 nm, 2.1 W/cm^2^).

**Figure S20.**
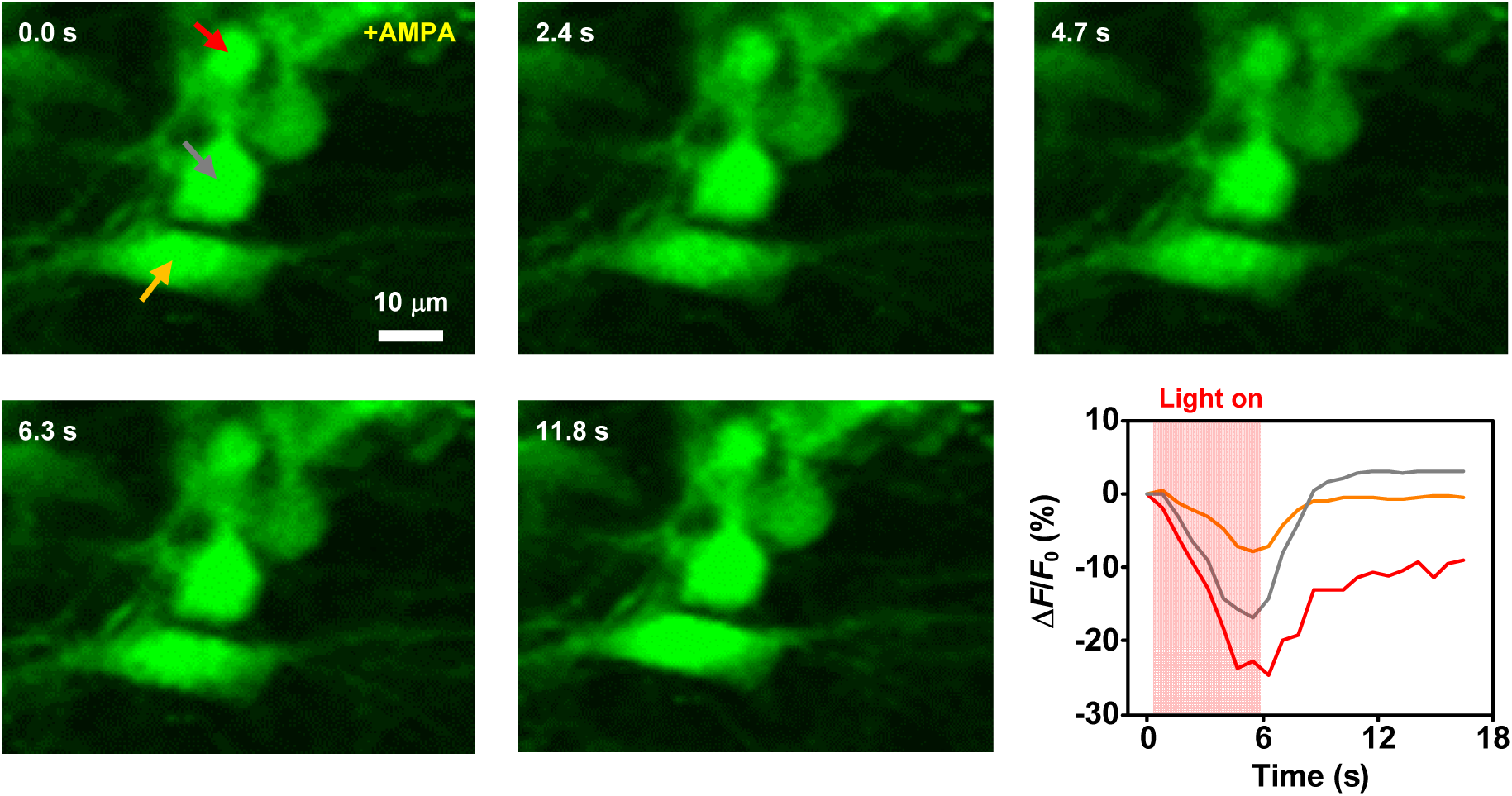
Calcium imaging of glial cells on n^+^p Si films. Glial cells can be optically inhibited to trigger intracellular calcium descend (green) over time. Arrows present the glial cells. The cells are initially activated by AMPA. The light is on from 0 s to 5 s (630 nm, 1.5 W/cm^2^).

**Figure S21.**
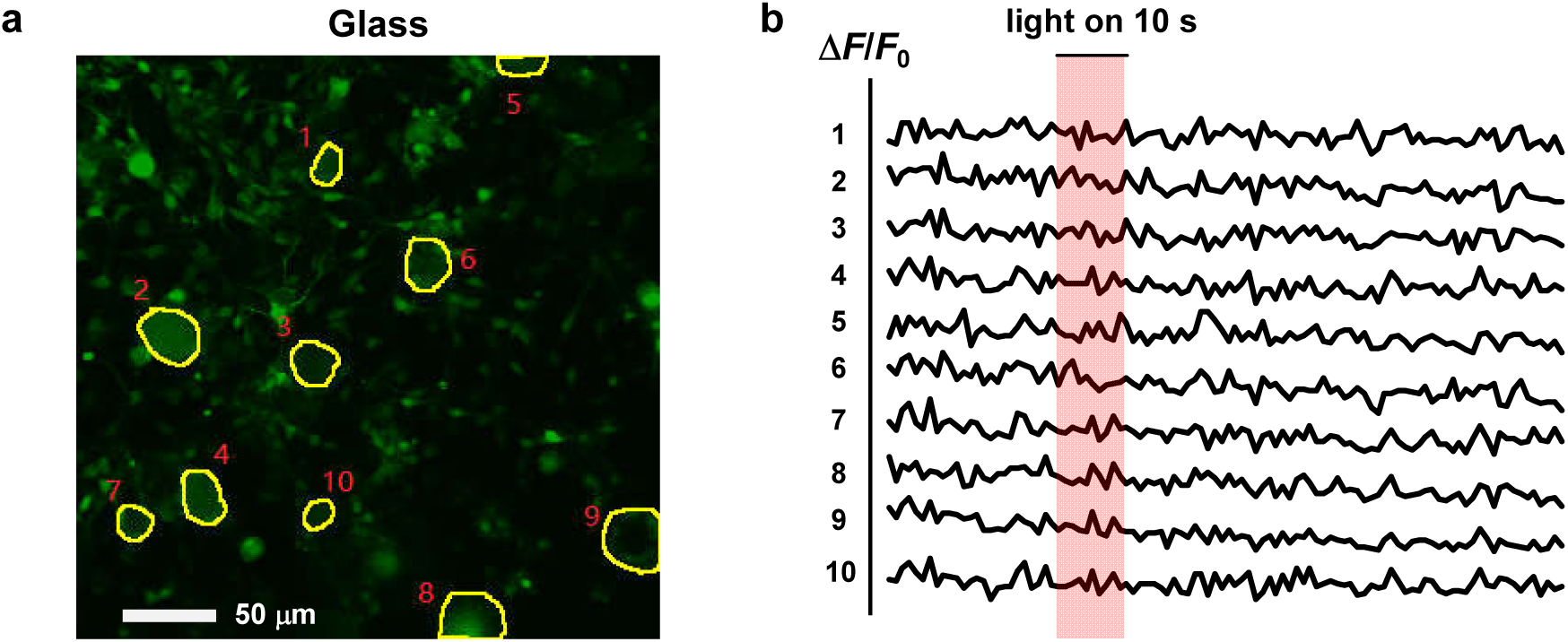
Calcium dynamics for neurons cultured on glass, as a control group. (a) Image of calcium loaded neurons cultured on glass (calcium, green; DRG neurons, yellow marked circles) from one region. (b) Calcium signal traces (Δ*F*/*F*_0_) of the marked DRG neurons from the region. Light duration 10 s, intensity 3.2 W/cm^2^. No obvious calcium fluorescence change is observed under illumination. Statistics are from 4 regions of 2 different cultures (total *n* > 30 neurons). All data are presented as means ± s.e.m.

**Figure S22.**
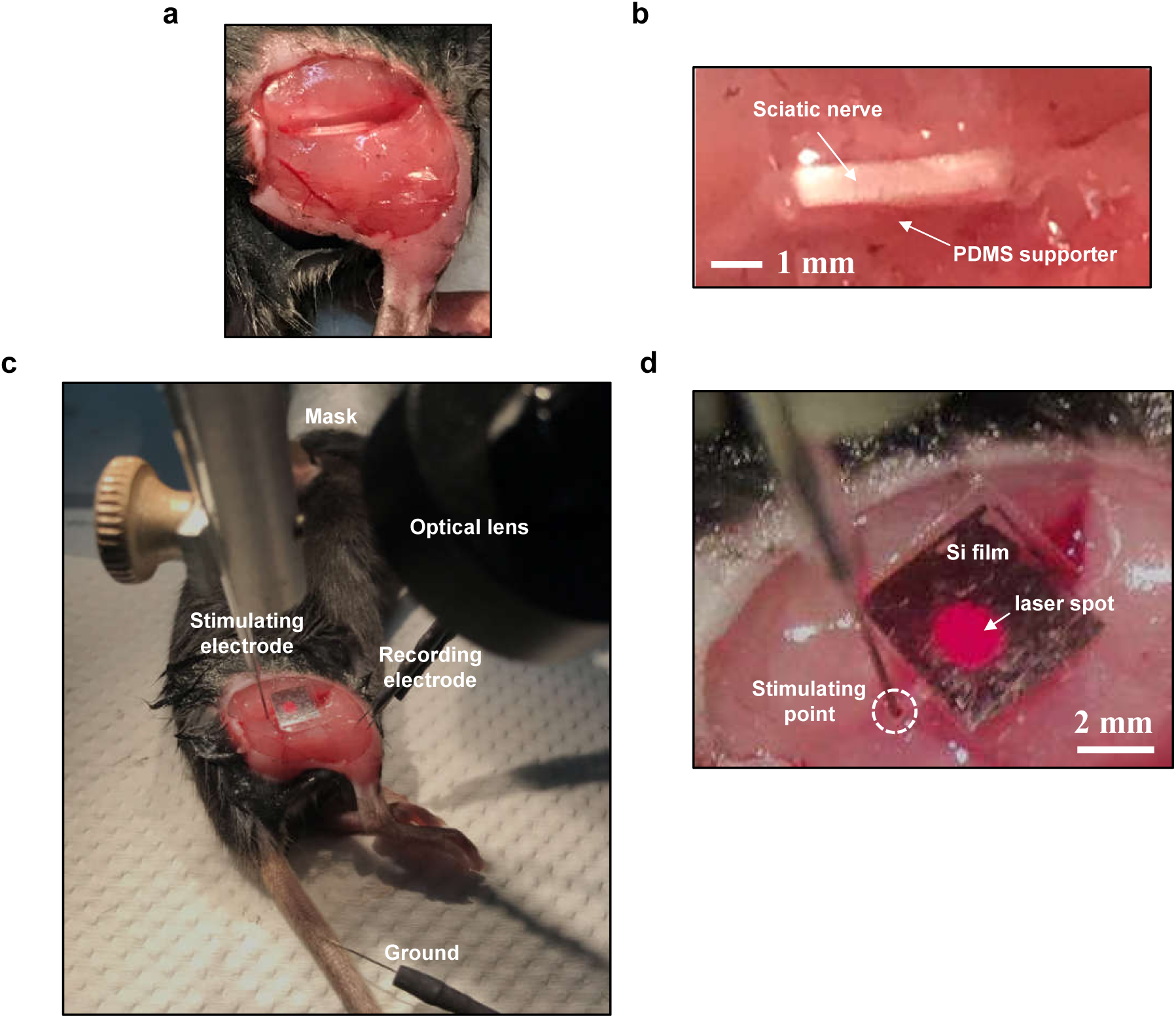
Scheme of the setup for modulating mice’s hindlimb movement. (a) The operation for exposing the sciatic nerve by tearing the connective tissue between the muscles without damage. (b) Fixing the sciatic nerve by a PDMS supporter. (c) The setup for recording and stimulation. (d) External metal electrode stimulating at the proximal position of nerve for evoking the CMAPs in the case of inhibition group. The laser spot illuminates the Si film mounted on the nerve.

**Figure S23.**
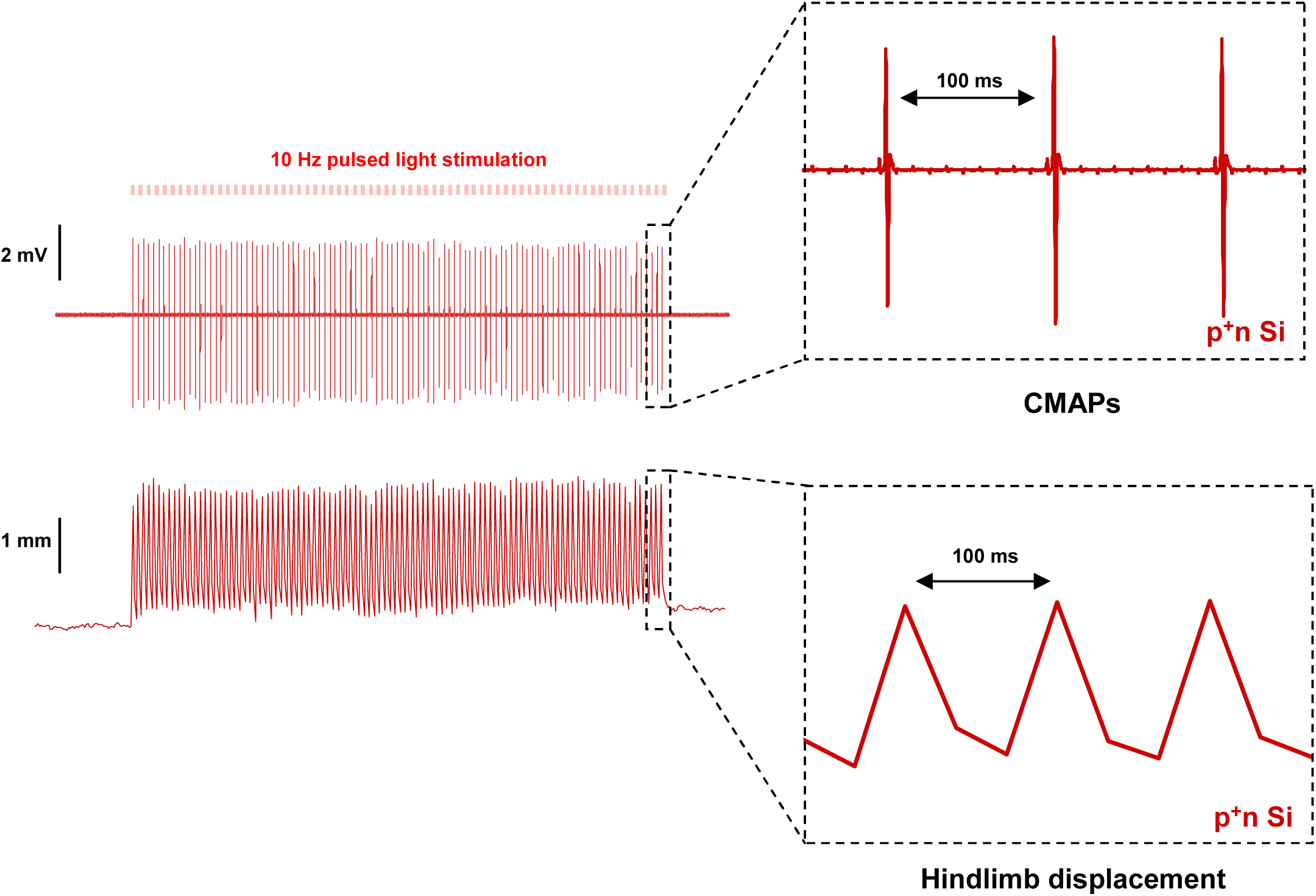
Example traces of recorded CMAPs (top) and displacements (bottom) evoked by p^+^n Si film under optical stimulation (10 Hz, 1-ms pulse, 0.9 W/cm^2^).

**Figure S24.**
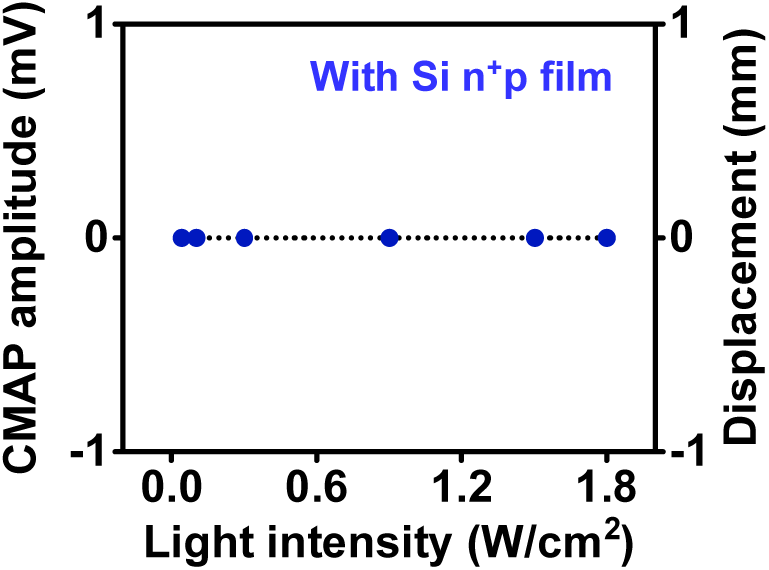
n^+^p Si film does not elicit hindlimb activities under optical stimulation (0.4 Hz, 1-ms pulse). Pulses are repeated 9 times on two mice.

**Figure S25.**
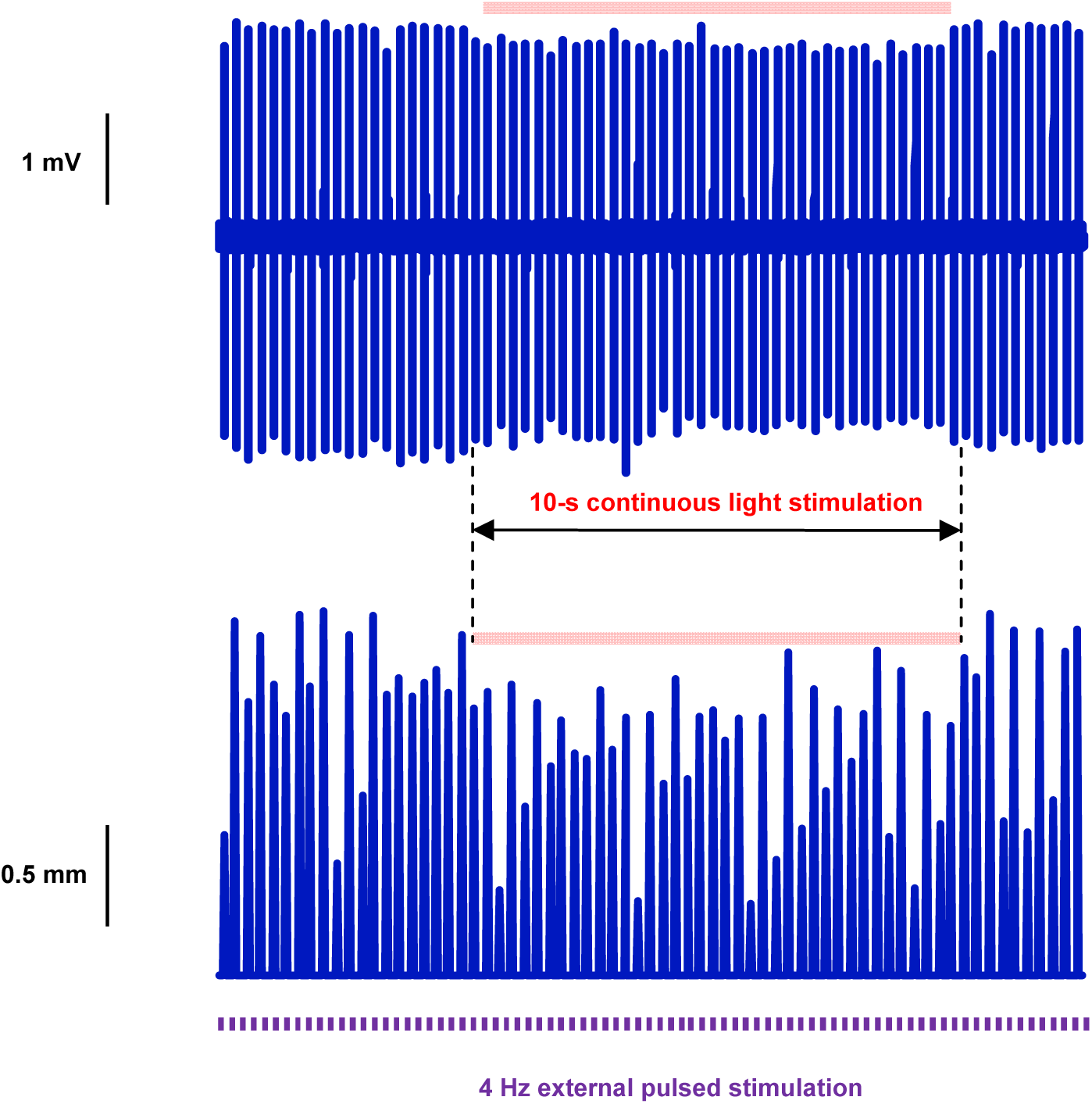
Example traces of decreased hindlimb movements (top: CMAPs; bottom: displacements) caused by n^+^p Si film under optical stimulation (10-s continuous stimulation, 0.9 W/cm^2^). The hindlimb activity is initially evoked by injecting pulsed current from external stimulation electrode (1-ms pulse, 4 Hz).

**Figure S26.**
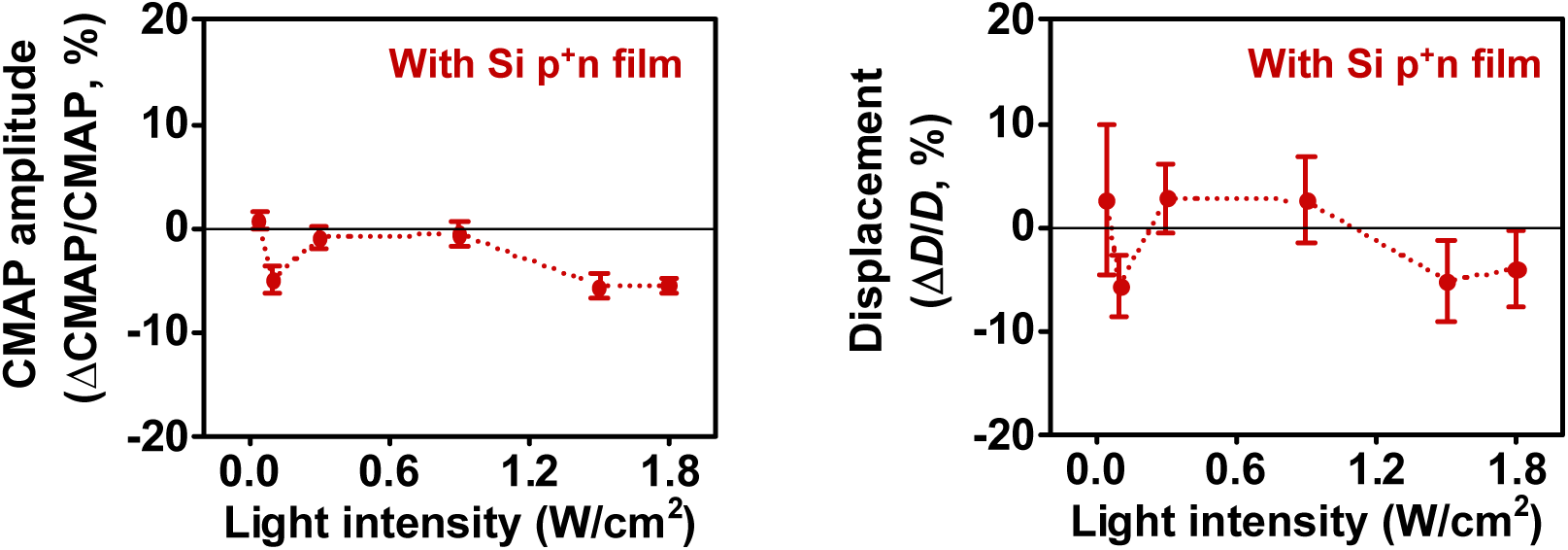
Statistics of hindlimb movements (Left: CMAP amplitude; Right: displacement) by p^+^n Si film under at varying light intensities (10-s continuous light stimulation), showing little inhibition effects. The hindlimb activity is initially evoked by injecting pulsed current from external stimulation electrode (1-ms pulse, 4 Hz). Each light intensity repeated more than 2 times and each time inhibited 40 trails (40-time hindlimb movement) from three mice. Data are presented as means ± s.e.m.

**Figure S27.**
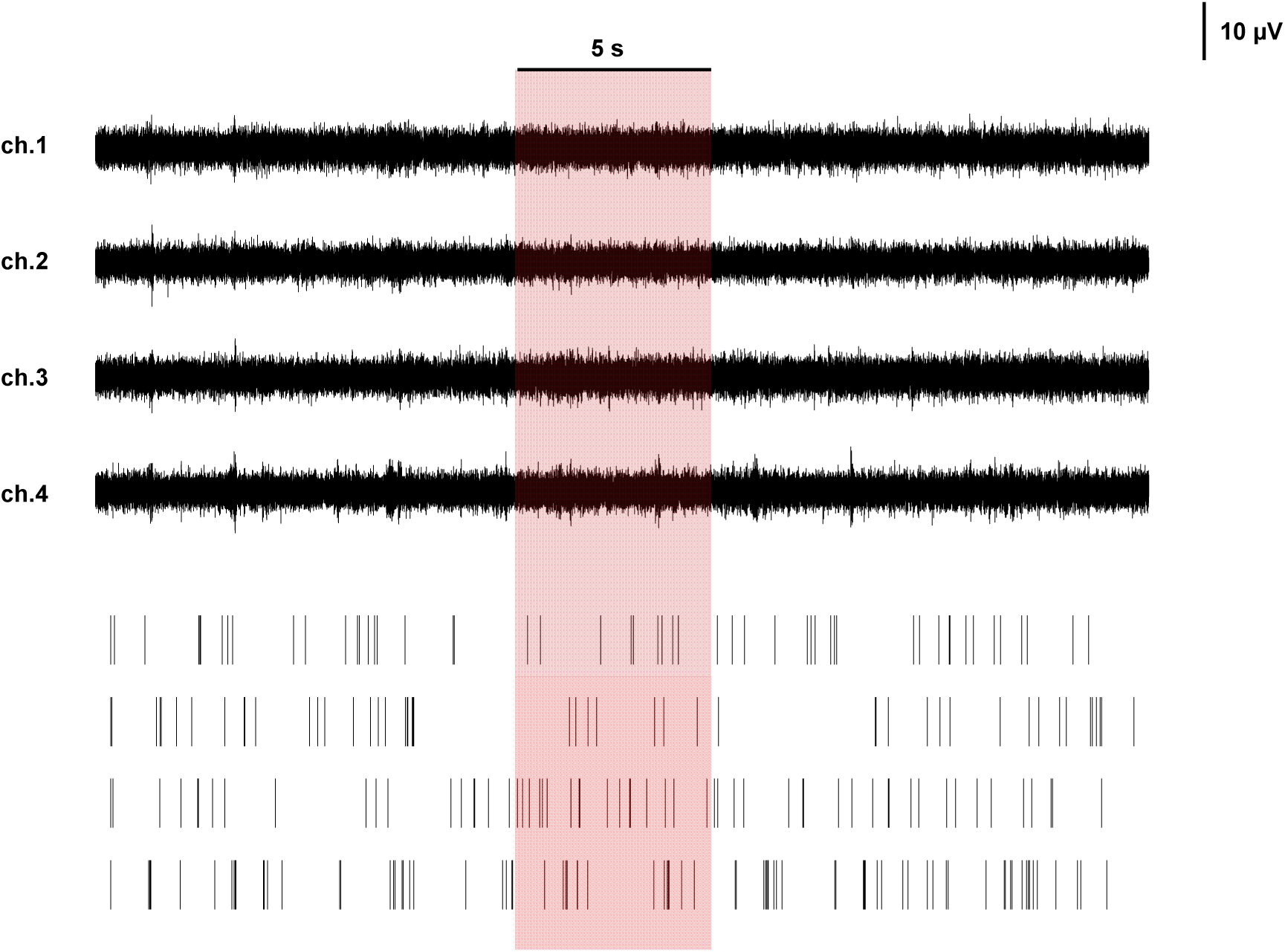
Example traces of neural response raw data (top) and corresponding spike raster plots (bottom) extracted from four channels in a single trial of optical stimulation on p-Si (0.06 W/cm^2^, 5-s duration), which does not evoke obvious increased spike frequencies during and after stimulation.

**Figure S28.**
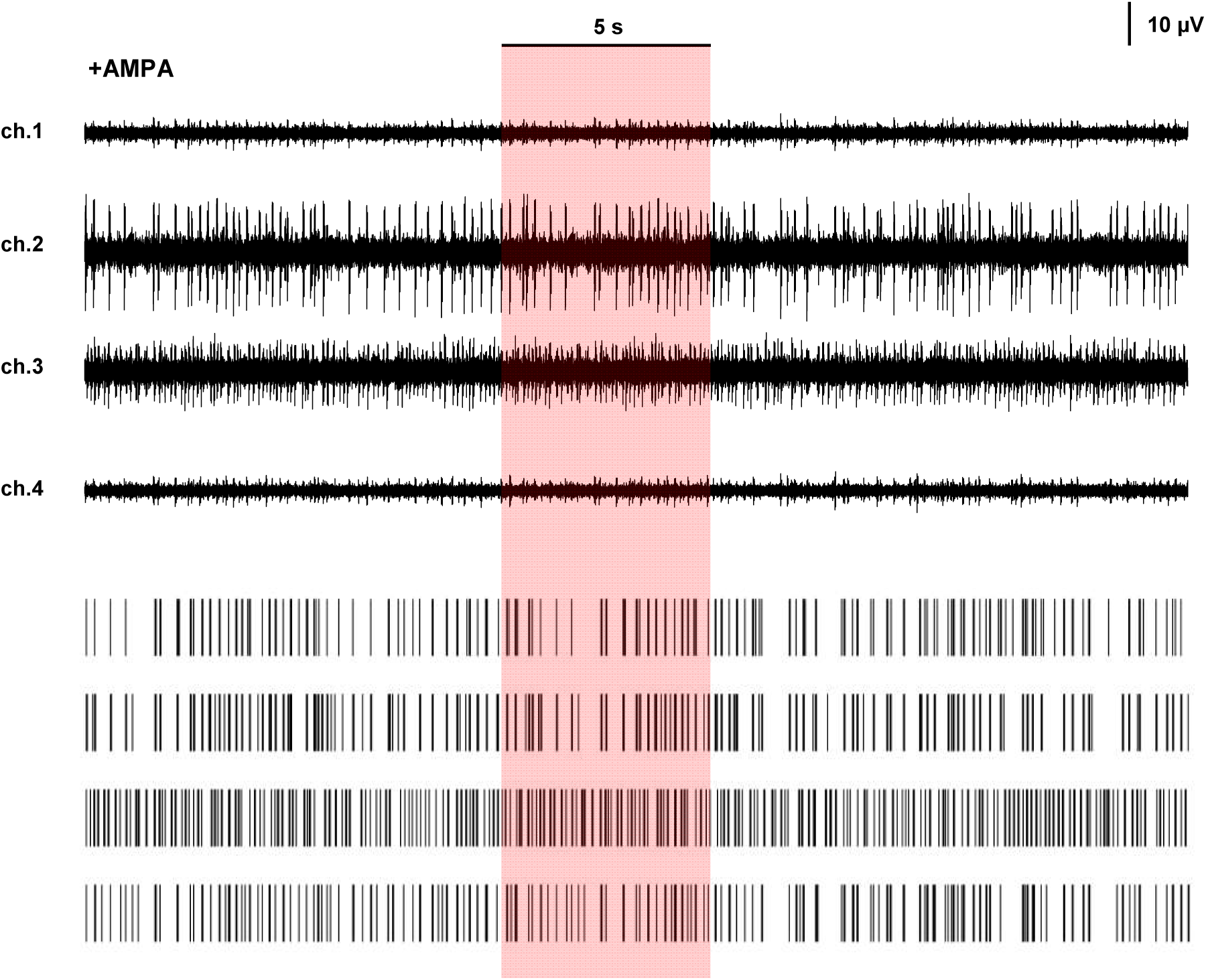
Example traces of neural response raw data (top) and the corresponding spike raster plots (bottom) extracted from four channels in a single trial of optical stimulation on p-Si (0.03 W/cm^2^, 5-s duration), which does not evoke obvious decreased spike frequencies during and after stimulation. AMPA is initially applied to activate the neural activity.

**Figure S29.**
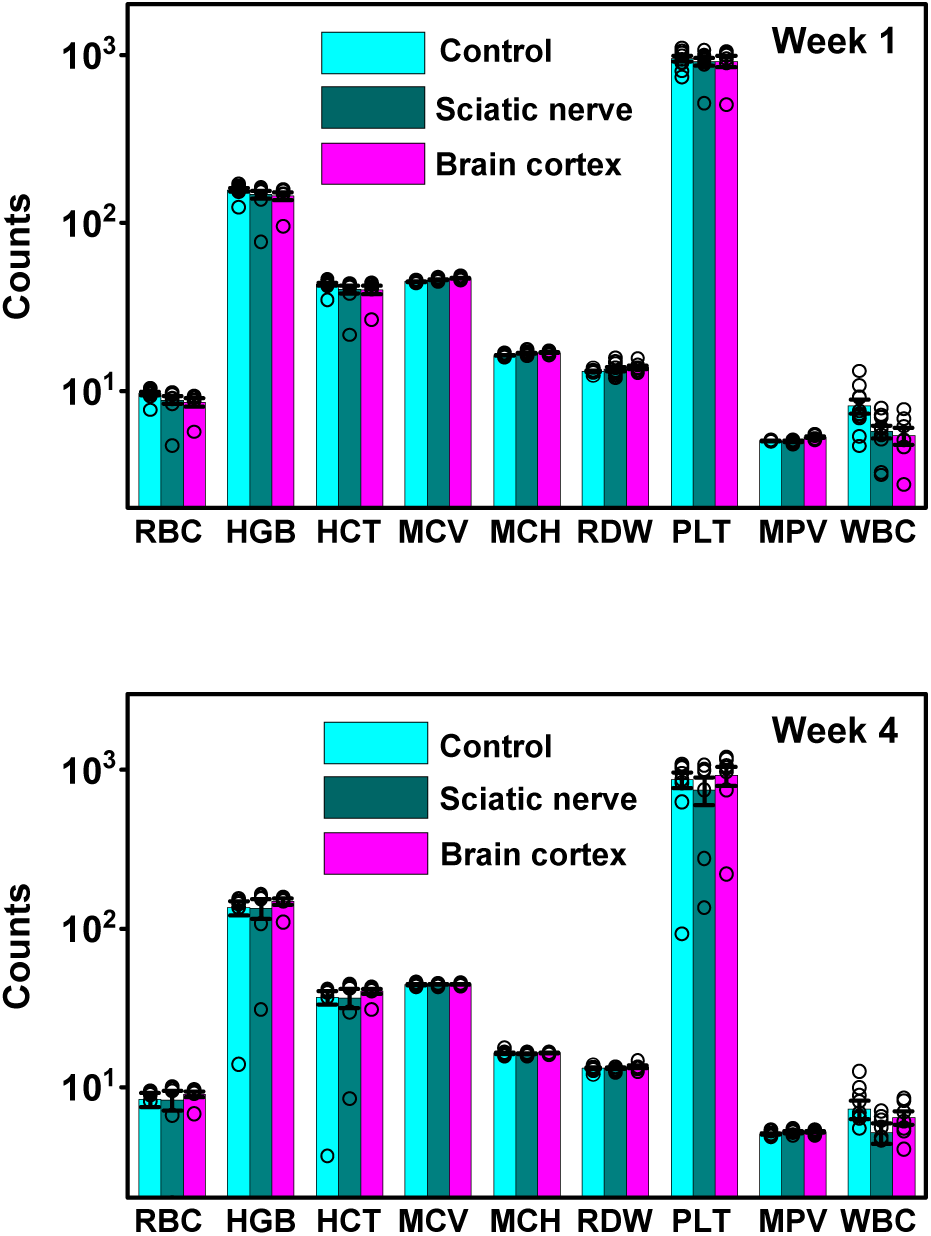
Analysis of complete blood counts for the mice with Si films wrapped around the sciatic nerve or mounted on the brain cortex, 1 and 4 weeks post implantation. The control group has no implants. *n* = 5 mice for each case. RBC, red blood cell (×1,000,000 µl^−1^); HGB, blood haemoglobin level (gl^−1^); HCT, haematocrit level (%); MCV, mean corpuscular volume (fl); MCH, mean corpuscular haemoglobin (pg); RDW, red cell distribution width (%); PLT, platelet count in blood (×1,000 µl^−1^); MPV, mean platelet volume (fl); WBC, white blood cell (×1,000 µl^−1^). Data are presented as means ± s.e.m.

**Figure S30.**
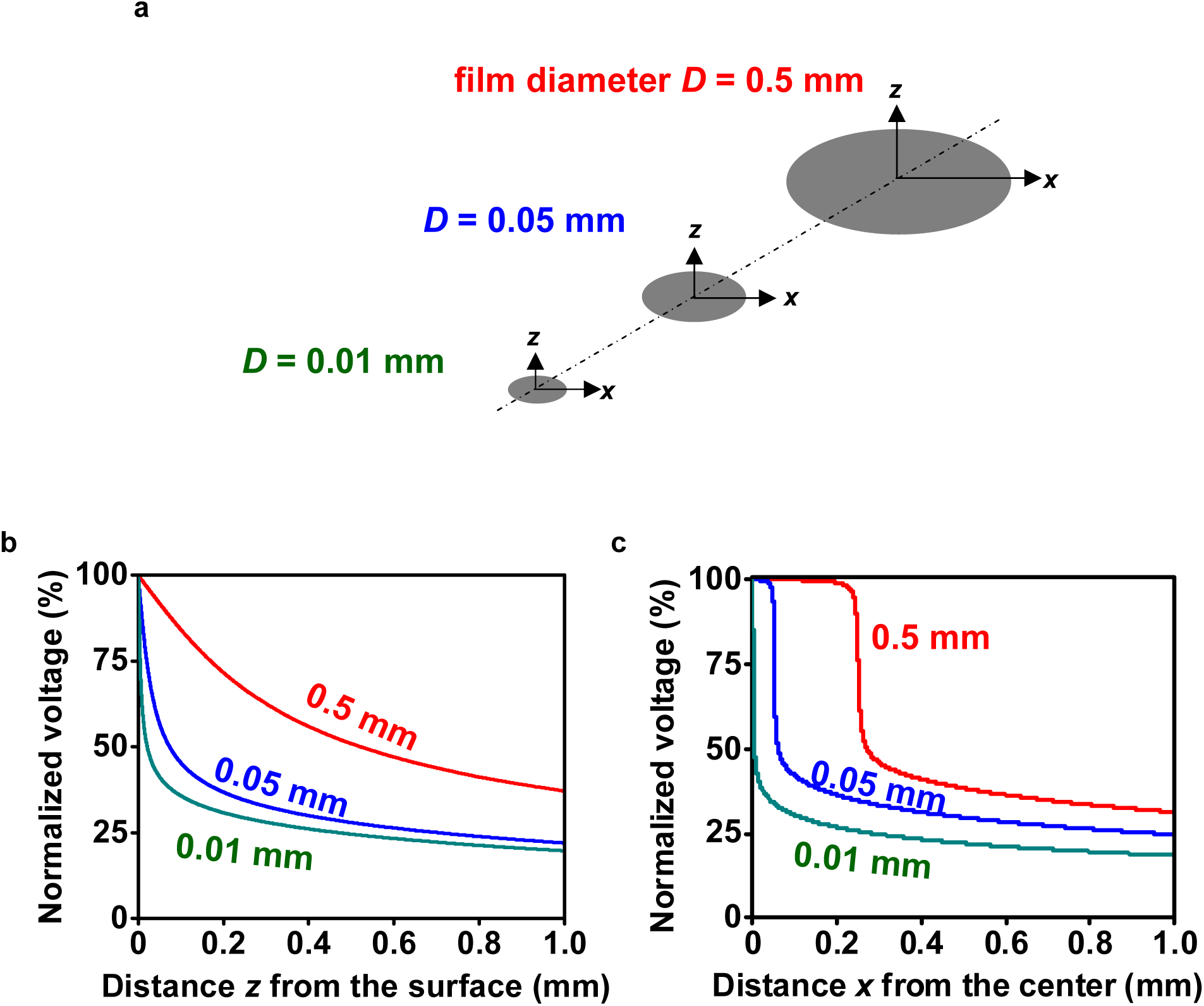
Simulated photovoltage distributions as a function of the Si film size. (a) Scheme of the setup for Si films with different sizes (diameters of 0.5 mm, 0.05 mm and 0.01 mm). (b, c) Simulated photovoltage at a function of (b) the vertical distance *z* and (c) the lateral distance *x* from the center of the film. Here we assume the illuminated spot has a fixed diameter of 2 mm. The photogenerated voltages reach ∼50% of the maximum values at *z* = 498 µm, 72 µm, 20 µm or *x* = 271 µm, 62 µm, 6 µm for film diameters of 0.5 mm, 0.05 mm and 0.01 mm, respectively.

**Movie S1.**
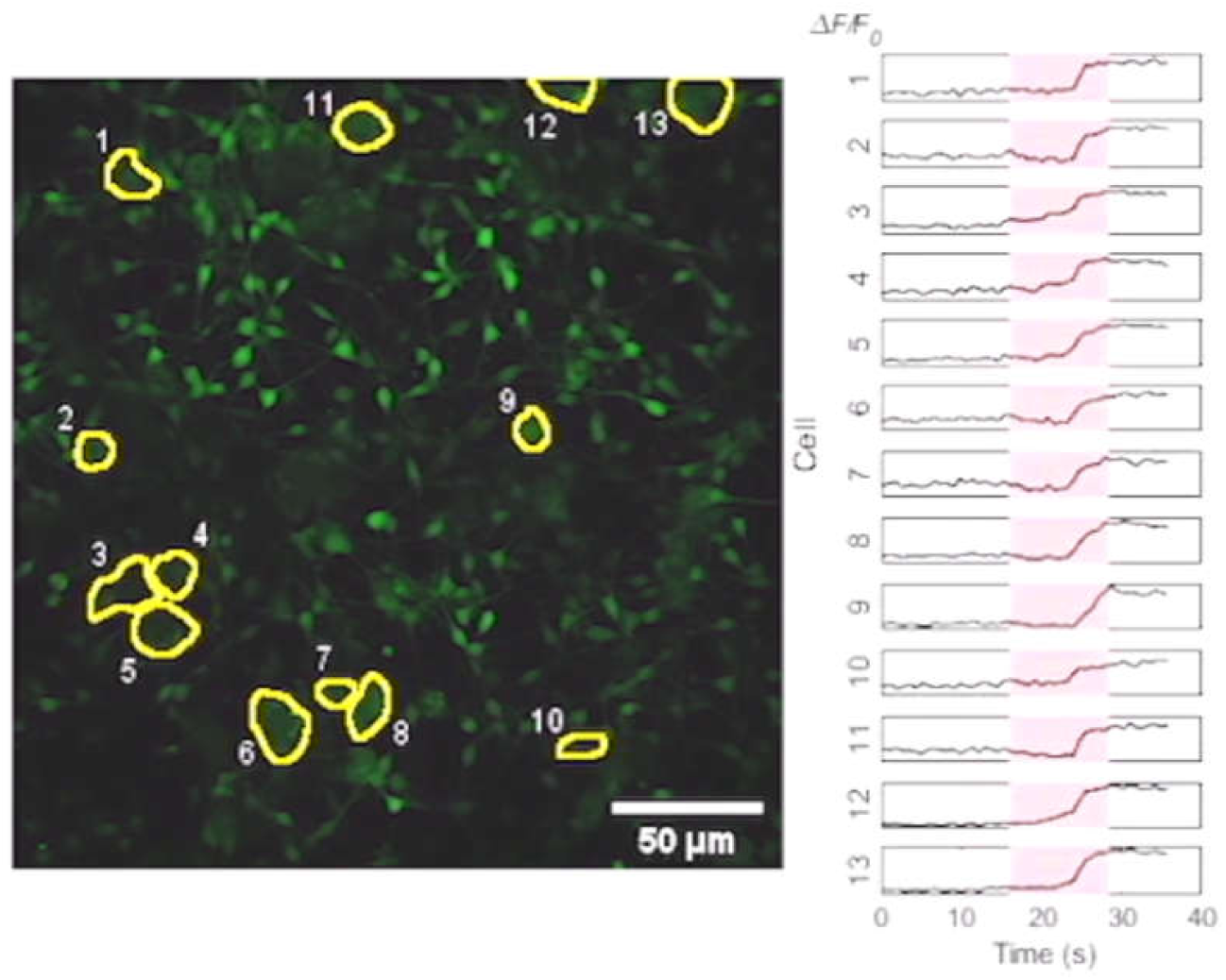
Video showing increased Ca^2+^ fluorescence in cultured DRGs evoked by a p^+^n Si film under optical stimulation.

**Movie S2.**
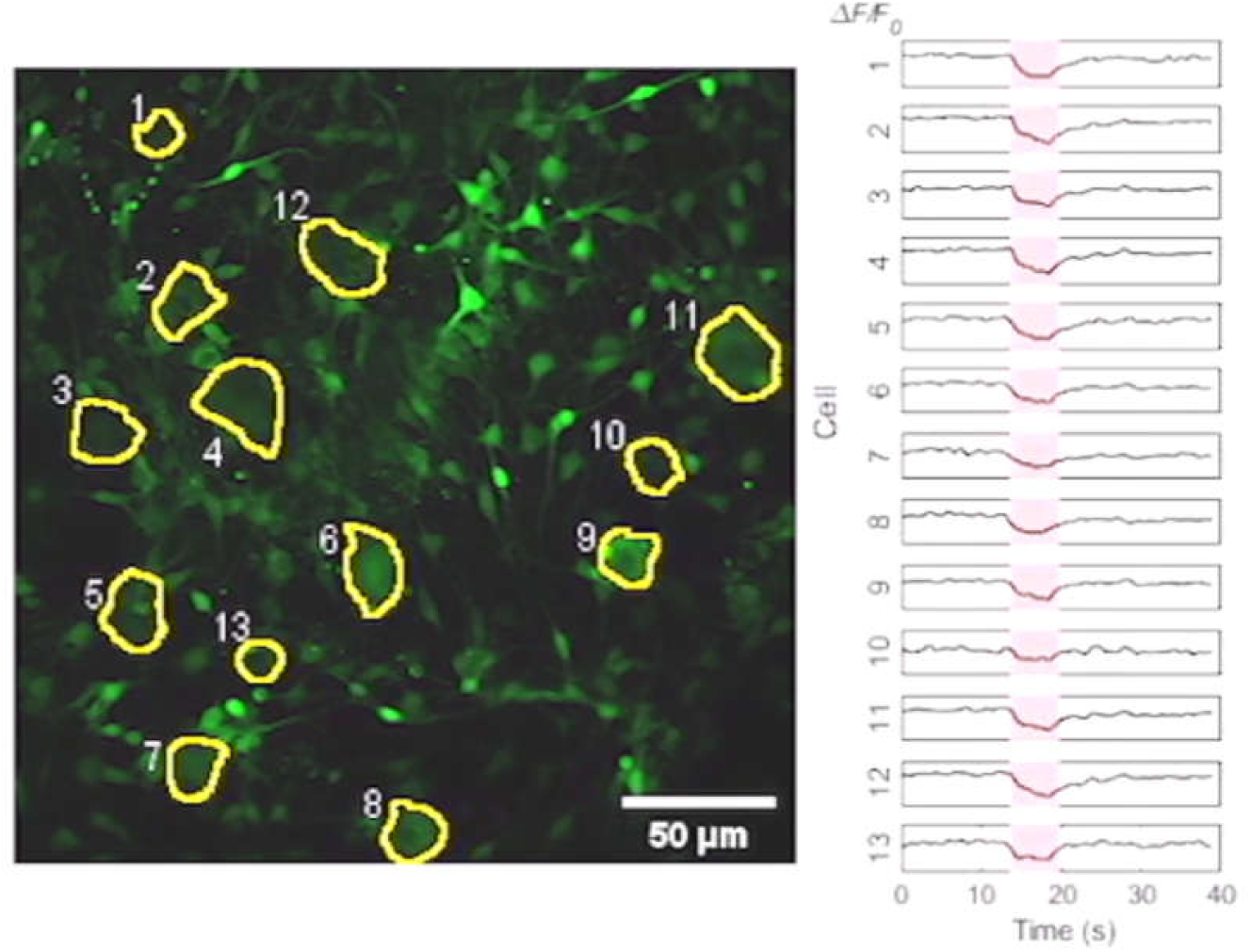
Video showing decreased Ca^2+^ fluorescence in cultured DRGs suppressed by an n^+^p Si film under optical stimulation. AMPA is initially applied for cell activation.

**Movie S3.**
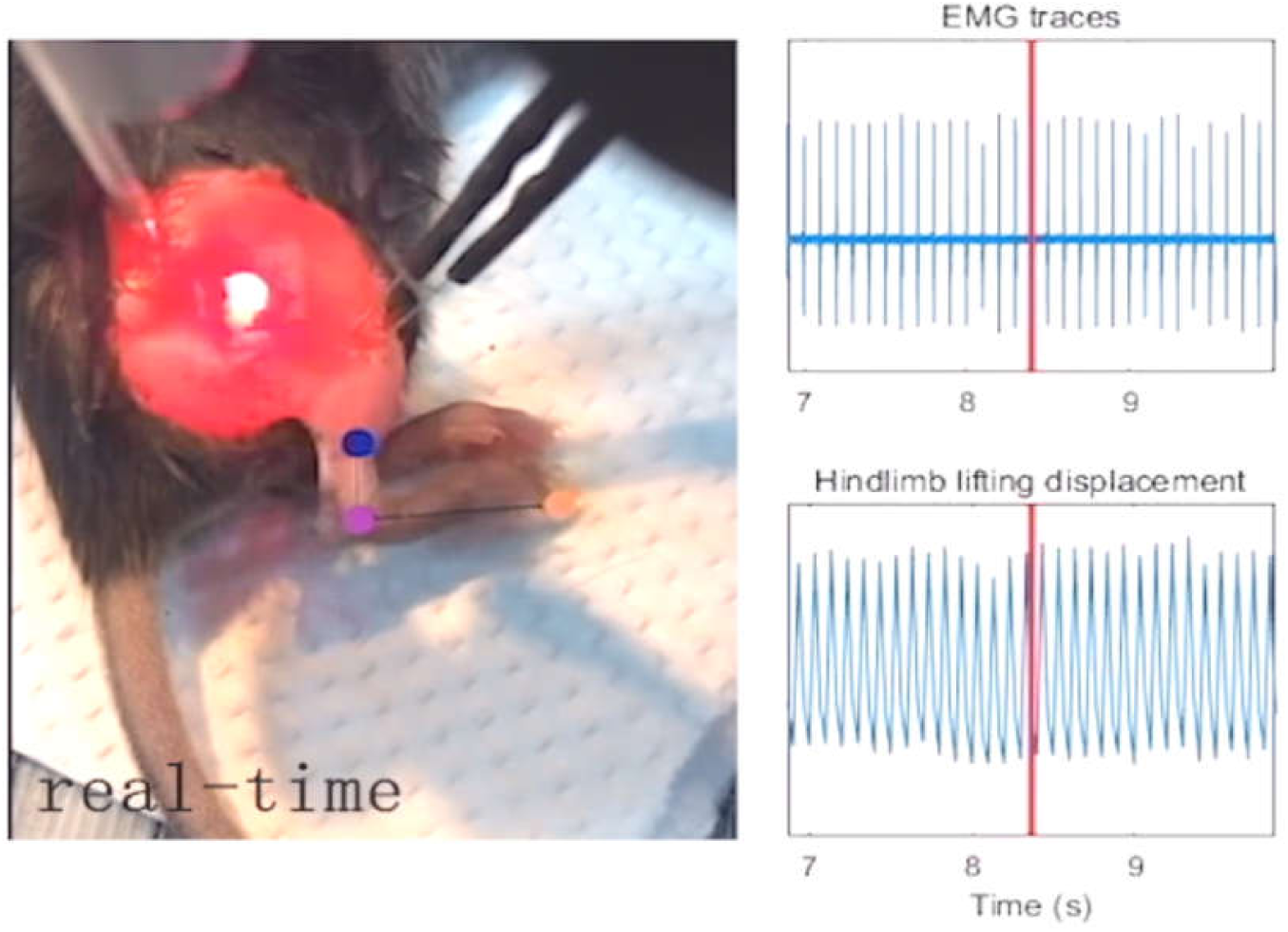
Video showing evoked CMAPs and hindlimb lifting by exciting in the sciatic nerve with a p^+^n Si film under optical stimulation.

**Movie S4.**
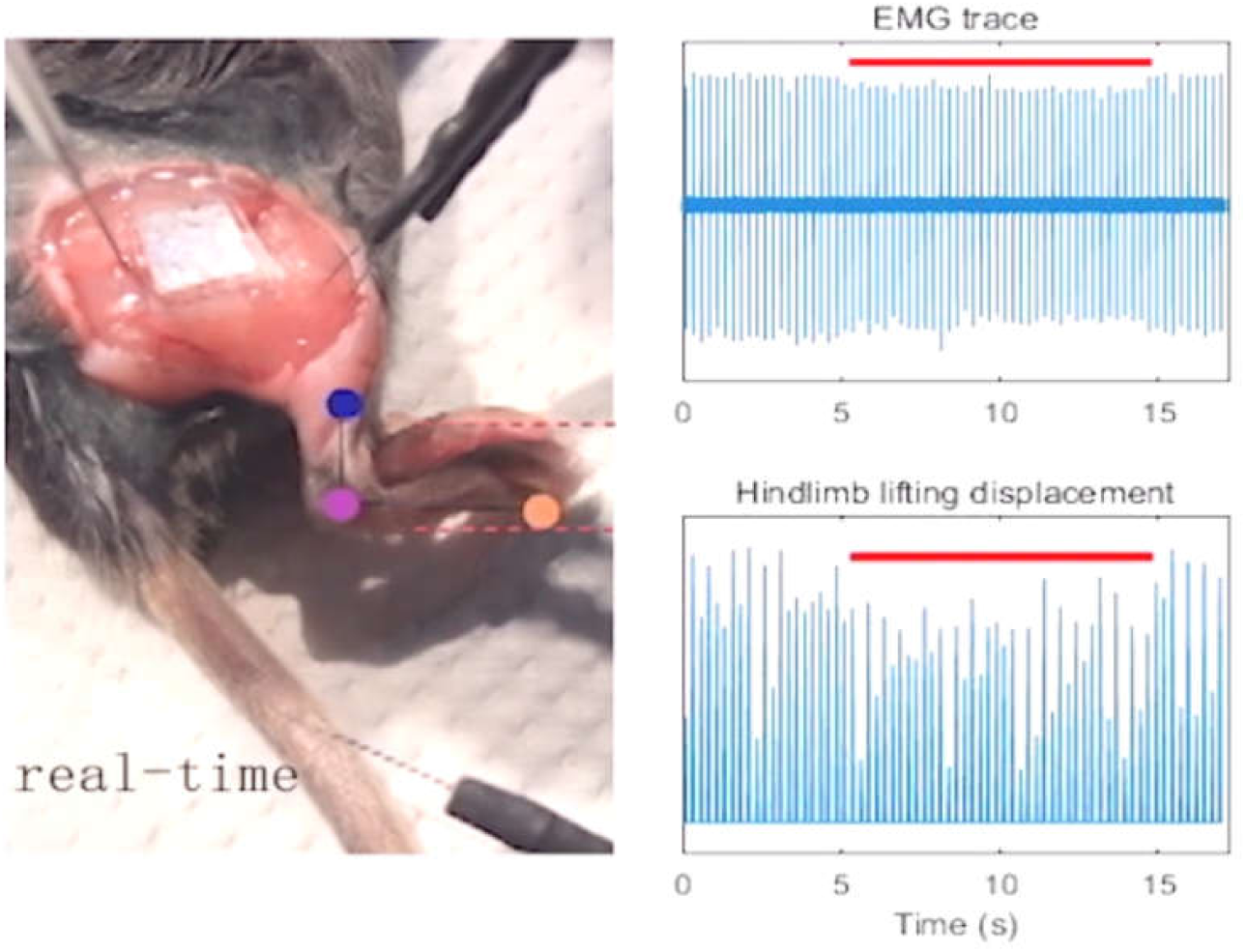
Video showing decreased CMAPs and hindlimb lifting by inhibiting in the sciatic nerve with an n^+^p Si film under optical stimulation.

